# *Ocimum sanctum, OscWRKY1*, regulates phenylpropanoid pathway genes and promotes resistance to pathogen infection in Arabidopsis

**DOI:** 10.1101/2022.01.12.474522

**Authors:** Ashutosh Joshi, Gajendra Singh Jeena, Shikha, Ravi Kumar, Alok Pandey, Rakesh Kumar Shukla

## Abstract

WRKY transcription factor (TF) family regulates various developmental and physiological functions in plants. *PAL* genes encode enzymes which are involved in plant defense responses, but the direct regulation of *PAL* genes and phenylpropanoid pathway through WRKY TF’s is not well characterized. In the present study, we have characterized an *OscWRKY1* gene from *O. sanctum* which shows induced expression after methyl jasmonate (MeJA), salicylic acid (SA), and wounding. Recombinant OscWRKY1 protein binds to the W-box *cis*-element TTGAC[C/T] and activates the reporter gene in yeast. Overexpression of *OscWRKY1* enhances Arabidopsis resistance towards *Pseudomonas syringae pv. tomato Pst* DC3000. Upstream activator sequences of *PAL* and *C4H* have identified the conserved W-box *cis*-element (TTGACC) in both *O. sanctum* and Arabidopsis. OscWRKY1 was found to interact with W-box *cis*-element present in the *PAL* and *C4H* promoters. Silencing of *OscWRKY1* using VIGS resulted in reduced expression of *PAL, C4H, COMT, F5H* and *4CL* transcripts. *OscWRKY1* silenced plants exhibit reduced PAL activity, whereas, the overexpression lines of *OscWRKY1* in Arabidopsis exhibit increased PAL activity. These results revealed that *OscWRKY1* positively regulates the phenylpropanoid pathway genes and enhances the resistance against bacterial pathogen in Arabidopsis.

## Introduction

*O. sanctum,* commonly referred to as “holy basil,” is a member of the Lamiaceae family. It has been significantly used worldwide due to its immense pharmacological properties and economically important aromatic oils related to terpenoid biosynthesis (Rastogi et al., 2014). *O. sanctum* leaves contains many biologically active compounds like triterpenoids, flavonoids, saponins and tannins (Jaggi et al., 2003). *O. sanctum* has antihistaminic, larvicidal, antibacterial, radioprotective, cardio-protective, anti-genotoxic, neuro-protective, anti-anaphylactic, wound healing, and antidiabetic activity (Rahman et al., 2011).

TFs usually interacts with a *cis*-element in the promoter of downstream genes, hence providing the regulation of the signaling cascade. (Agarwal and Jha, 2010). Generally, a plant-specific WRKY TF family is mainly characterized by the presence of a highly conserved WRKYGQK peptide sequence and a C2H2 or C2HC zinc finger motif (Eulgem et al., 2000). These regulatory TFs preferentially bind with the W box *cis*-elements (T/C-TGAC-T/C) present in their promoter, thereby differentially regulating the expression of target genes. Activation or repression of target genes is mainly regulated at transcriptional, translational, and domain levels through W-box consensus sequences (Phukan et al., 2016).

Based on the number of WRKY domains present, WRKY proteins are categorised into three major types. Group-I WRKY proteins contain two WRKY domains, whereas Group-II and -III WRKY proteins include a single WRKY domain. A zinc finger motif of the C2H2 type (C-X4-5-C-X22-23-H-X1-H) is found in Group-I and Group-II WRKY proteins, whereas the C2-HC type (C-X7-C-X23-H-X1-C) is found in Group-III WRKY proteins. Based on their primary amino acid sequence, Group-II WRKY proteins are classified into different subgroups IIa, IIb, IIc, IId, and IIe (Rushton et al., 2010). Group-I WRKY proteins feature functionally distinct WRKY domains, and it was discovered that the C-terminal domain of proteins binds to target DNA in a sequence-specific manner (Eulgem et al., 2000). The WRKY TF family is well known for its important function in controlling plant biotic and abiotic stress tolerance. WRKY members regulate the genes related to metabolite synthesis, hence regulating the production of valuable natural products. They are found to regulate the three major classes of plant secondary metabolite biosynthetic pathways *i.e.,* phenylpropanoids, alkaloids, and terpenoidal biosynthesis (Schluttenhofer and Yuan, 2014; Mishra et al., 2013). It has been reported that several *WRKY* genes were induced during pathogen infection, elicitor treatment, or hormonal challenges in several plant species (Dong et al., 2003). *AtWRKY70* in Arabidopsis is a key player in the defense signaling pathways mediated by jasmonic acid and SA, which leads to the enhanced pathogen resistance against specific bacteria and fungi (Li et al., 2004). Similarly, *AtWRKY33* confers increased pathogen resistance against necrotrophic *Botrytis cinerea via* activation of tryptophan-derived camalexin biosynthesis (Mao et al., 2010). A WRKY TF from potato, *StWRKY1* was involved in the regulation of phenylpropanoid biosynthesis against infection of *Phytophthora infestans* (Yogendra et al., 2015).

Phenylalanine gives rise to cinnamic acid, which acts as an intermediate in the formation of phenylpropanoids. The non-oxidative deamination of phenylalanine results in the formation of *trans*-cinnamate, which is catalyzed by PAL (Phenylalanine ammonia-lyase) enzyme. This is an important regulatory step between primary and secondary metabolism (Huang et al., 2010; Vogt T, 2010). PAL plays a crucial role in plant defense mediated *via* the biosynthesis of SA, which acts as an essential signaling molecule in plant systemic resistance (Chaman et al., 2003; Nugroho et al., 2002). Expression studies of *PAL* reveal its corresponding response to various abiotic and biotic stresses like UV irradiation, nutrient depletion, wounding, extreme temperatures, and pathogen infection (Jin et al., 2013; Payyavula et al., 2012; Huang et al., 2010; Shine et al., 2016). Another important enzyme of the phenylpropanoid biosynthetic pathway is C4H, which is involved in synthesizing the building blocks of the lignin polymer. C4H mainly converts the *trans*-cinnamic acid into *p*-coumaric acid, which is the first hydroxylation step in the biosynthesis of lignin, hydroxycinnamic acid esters, and flavonoids (Blount et al., 2000; Blee et al., 2001).

In the present manuscript, our study elucidated the functional role of *OscWRKY1* TF from *O. sanctum* in directly regulating the phenylpropanoid pathway. The interaction of *OscWRKY1* with *PAL* and *C4H* promoters in both *O. sanctum* and Arabidopsis suggest the possible mechanism of *OscWRKY1* in regulating the phenylpropanoid pathway leading to the enhanced *Pst* DC3000 resistance in the OE lines of Arabidopsis. We have utilized the VIGS for the first time in *Ocimum sanctum* to functionally characterize the *OscWRKY1*. *OscWRKY1* silencing provides a novel molecular insight into its role in directly regulating the phenylpropanoid pathway genes in *O. sanctum.*

## Materials and methods

### Identification of some putative *WRKY* transcripts from *O. sanctum*

The various WRKY protein sequences were identified using *O. sanctum* transcriptome data (SRA Study accession number SRP039008). Full-length WRKY protein sequences in *Arabidopsis thaliana* and *Oryza sativa* (http://www.arabidopsis.org/) were used as reference sequences. The BLASTX search tool was used to manually validate selected WRKY proteins. All WRKY sequences, whether complete or partial, were included in this quest. From all these WRKY sequences full-length WRKY protein sequences were selected and grouped into WRKY subgroup I, II, and III, according to the classification data available from TAIR (Arabidopsis) WRKY TF database.

### Phylogenetic analysis based on conserved WRKY Domains

CLUSTALW performed multiple sequence alignment of the amino acid sequences of both full-length and partial WRKY sequences. MEGAX with pairwise distances and the neighbor-joining algorithm was used to evaluate the phylogenetic and evolutionary relationships between various WRKYs. Only protein sequences having the highest homology with OScWRKY1 sequence were used in the multiple sequence alignment and phylogenetic study.

### Plant Growth and treatments

*O. Sanctum* two-month-old plants (var: CIM Ayu) were grown and collected from CSIR-CIMAP, hariti growth chamber. Plants were sprayed with a solution of 200 μM MeJA and SA in dimethyl sulfoxide (DMSO) and Triton-X. In control plants, only DMSO and Triton-X-containing solutions were sprayed. To maintain the proper transpiration rate, perforated plastic bags were used to cover the samples. At different time intervals, samples treated with MeJA/SA were collected and thoroughly washed with autoclaved water to eliminate contaminants. To avoid substantial injury, the whole plant was perforated with sterile needles to treat wounds in the *O. sanctum.* For further analysis, samples were taken from treated and control plants at 1, 3, and 5-hour time intervals. Arabidopsis (WT Col-0) and transgenic *OscWRKY1* seeds were sown in a soilrite mixture and maintained in the growth chamber at 22°C, 16:8 light: dark, and 65 % relative humidity.

### Total RNA isolation, cDNA synthesis, and qRT-PCR

The leaf and root samples were rinsed twice with DEPC treated water before being crushed in liquid N2. The NucleoSpin RNA kit (Macherey Nagel) was utilized to isolate RNA from plant samples. The quality of total RNA was monitored by using Nanodrop and running on 2% agarose gel. First-strand cDNA was synthesized from 1μg of total RNA utilizing a high-efficiency cDNA reverse transcription kit (Thermo Scientific). Total cDNA products were quantified and checked for purity. The qRT-PCR reactions were set up for 20 μL, each containing 100 ng of total cDNA, 0.4 μM of forward and reverse primers, 10 μL of SYBR Premix Ex Taq^TM^ II (Takara), and 0.4μL of reference dye. The amplification conditions for all the primer pairs were: 95*°C* initial denaturation for 10 min; 95°C denaturation for 15 seconds, 60*°C* primer annealing/elongation for 1 minute (40 cycles). The fluorescence was detected and measured during the last step of each cycle.

After each PCR loop, a melting curve analysis was performed by progressively rising the fluorescence temperature from 60 to 95°C. The qRT-PCR experiments used the 7500 FAST Real-Time PCR system (Applied Biosystems, USA) (Fig. S10). Actin and ubiquitin were used to normalize gene expression as endogenous controls. To evaluate relative transcript expression, the 2^-ΔΔCt^ method was used. Both tests were carried out with three biological and experimental replicates, and the data were then statistically analyzed. Real-time PCR primers were produced using the unique region of the transcript in table S1.

### Electrophoretic mobility shift assay (EMSA) and transactivation β-galactosidase assay

The *O. sanctum* transcriptome data set was used to select *OscWRKY1,* which was then cloned into the bacterial expression vector pGEX4T2 using *EcoRI* and *XhoI* restriction sites. The construct was confirmed using digestion and sequencing. The reading frame was retained in addition to GST. After that, the *OscWRKY1-GST* cloned plasmid was transformed in BL21 (CodonPlus, DE3 strain), and then induced with IPTG (0.4 mM). The recombinant OscWRKY1 was affinity purified using GST beads (Amersham). The W-box sequence TTGAC[C/T] *cis*-element probes were developed, as were their mutated counterparts (TTGACG, TTCACT) (Table S1). The EMSA was carried out utilizing the manufacturer’s instructions (DIG-gel shift kit 2^nd^ generation kit, Roche).

*OscWRKY1* was cloned into the *Saccharomyces cerevisiae* expression vector pGBKT7 and fused to the GAL4-DNA-binding domain. Positive clones were confirmed using sequencing and restriction enzyme digestion *(NdeI* and *BamHI).* The positive cloned plasmid was transformed in yeast Y187 *(S. cerevisiae),* containing *LacZ* gene with modified GAL4 promoter, and the resulting colonies were selected in the dropout medium (SD-Trp-Ura). The selected positive colonies were further streaked on an SD-Trp-Ura replica plate for the positive transactivation assay. The blue color intensity was determined using X-gal as a substrate and a colony lift assay with galactosidase (Phukan et al., 2018).

### Yeast One-Hybrid (Y1H) Assay

We identified the upstream promoter sequences of *PAL* and *C4H* from the genome sequence of both *O. sanctum* and Arabidopsis. Promoters were amplified using specific forward and reverse primers from their respective genomic DNA. *OscWRKY1* was cloned into the yeast expression vector pGADT7 for the Y1H assay, and the promoters of *O. sanctum* and Arabidopsis (*PAL* and *C4H)* were cloned into the pHIS-2.0 vector. In the yeast *5. cerevisiae strain* Y187, both cloned plasmid DNA were co-transformed employing lithium acetate-mediated (LiAc) yeast transformation. Desired cloned plasmid DNA and excess carrier DNA was transformed into the yeast competent cells (Gietz et al., 1992). The positive interaction has been checked by plating it on SD-His-Leu media. Positive colonies were streaked in YPD and SD-His-Leu selection dropout media. All yeast experiments were carried out using the Yeast Protocols Handbook (Clontech).

### PAL activity Assay

By using 100mM phosphate buffer (pH 6.0) with 4mM dithiothreitol, 2mM EDTA, and 2% polyvinylpyrrolidone, total protein was isolated from the leaf samples of Arabidopsis. In the leaf extract, PAL activity was determined as described in the earlier protocol (Song and Wang 2011). The protein extract was incubated for 50 mins at 30°C containing 2 mL borate buffer (0.01M, pH 8.7) and L-phenylalanine (0.02M, 1ml). The absorbance of the sample was taken at 290 nm before and after the incubation. For blank control, a reaction mixture without substrate was used and the experiment was performed in biological replicates, and for each extract, triplicate assays were performed. The amount of PAL required for producing 1 M cinnamic acid in 1 sec is defined as one katal and expressed in terms of nkatal mg^-1^ of protein.

### Bacterial resistance assay

Fully developed rosette leaves of WT Col-0, *OscWRKY1-L1* and *OscWRKY1-L2* Arabidopsis plants were syringe-infiltrated with *Pseudomonas syringae* pv. *tomato* DC3000 (*Pst* DC3000) resuspended in 10 mM MgCl_2_. Disease phenotype was observed and photographed at regular time points. For the quantification of bacterial growth, the leaf discs of area (1 cm^2^) from each genotype were collected at 0 and 2 days post-infection (dpi) and grounded in 1000 μL MilliQ. The homogenate was then serial diluted and plated on LA medium containing 50 μg ml^-1^ rifampicin. The bacterial colony-forming unit (CFU) was counted and the bacterial population [Log (CFU/cm^2^)] are presented. Two independent experiments were conducted and three biological replicates were taken in each experiment.

### *In planta* pathogen colonization using confocal microscopy

For the *in planta* pathogen colonization, WT Col-0, *OscWRKY1-L1* and *OscWRKY1-L2* Arabidopsis plants were syringe-infiltrated with GFP-tagged *Pst* DC3000 strain. At 2 dpi, images were collected from the infected leaf discs using confocal laser scanning microscopy. GFP fluorescence (488 nm excitation/493-598 nm emission) and red chlorophyll fluorescence (633 nm excitation/647-721 nm emission) was acquired on a Zeiss LSM 880 confocal microscope. Image acquisition was performed using ZEN blue software (Carl Zeiss, Oberkochen, Germany).

### Electrolyte leakage measurement

Four leaf discs each of diameter 4 mm were excised from each WT Col-0, *OscWRKY1-L1* and *OscWRKY1-L2* plants infiltrated with *Pst* DC3000 at 2 dpi. The leaf discs (each tube containing four discs per genotype) were then gently agitated in deionized water for 3 hours at 28°C. The initial conductivity was determined with the help of a conductivity meter (HORIBA Scientific, F74BW). Then, these tubes were boiled at 121° C for 20 min to release the total electrolytes and the conductivity was again measured. The values of the initial electrical conductivity, and conductivity obtained after autoclaving were used to calculate the percentage of electrolyte leakage relative to total electrolytes in each genotype. Two independent experiments were conducted and two biological replicates were taken in each experiment.

### Virus-induced gene silencing (VIGS) in *O. sanctum*

The pTRV1 and pTRV2 vectors based on the *Tobacco rattle virus* (TRV) were used to conduct VIGS in *O. sanctum.* (Liu et al. 2002). The cDNA sequences of *OscPDS* and *OscWRKY1* were retrieved from the transcriptome resource of the *O. sanctum*. Specific primers were designed utilizing the conserved domains of *OscPDS* and *OscWRKY1*. Both fragments were amplified using PCR and cloned into the pTRV2 vector, which was validated using restriction digestion and sequencing. Cloned plasmids were then individually transformed into *Agrobacterium tumefaciens* strain (GV3101). Overnight grown *Agrobacterium* suspensions containing pTRV1 and pTRV2 (pTRV2-*OscPDS* and pTRV2-*OscWRKY1*) were centrifuged and re-suspended in infiltration buffer (10 mM MES pH5.6, 10 mM MgCl_2_, 200 μM acetosyringone). Vacuum-aided agroinfiltration (Misra et al., 2020; Deng et al., 2021; Wang et al., 2018; Zhang et al., 2017) was employed to perform VIGS. Seeds of *O. sanctum* were surface sterilized and sown in the nursery bed. The cotyledonary leaf stage plants were tested for agro-infiltration using vacuum. The *Agrobacterium* pTRV1 and pTRV2 (pTRV2-*OscPDS* and pTRV2-*OscWRKY1*) suspensions were mixed in similar concentrations to achieve the optimal optical density OD600. *Agrobacterium* suspension was generated by mixing *Agrobacterium* possessing pTRV1 and pTRV2 vectors, served as control whilst using the infiltration buffer for mock control*. Agrobacterium* density was optimized for efficient VIGS and so suspension at OD600 of 0.5, 1.0, and 1.5 were also considered. At least, 100 mL of the final suspension was used. Healthy cotyledonary leaf stage seedlings with uniform growth were utilized for the vacuum-aided infiltration method. The seedlings were uprooted, washed with Milli-Q, and were then inverted into the suspension solution in a desiccator jar connected to the vacuum pump. A vacuum was generated with the help of a pump to further generate 750 mm Hg pressure. The pressure was applied for an additional 90 s (Misra et al., 2020) and was released slowly causing infiltration of the *Agrobacterium* suspension into the immersed plant leaves. The seedlings designated to serve as a mock were also immersed in the infiltration buffer. The seedlings were removed and the excess buffer was drained in blotting paper. The seedlings were then transplanted into plastic pots containing soilrite mix, covered with PVC cling film to maintain an elevated humidity level, and left in the dark overnight at room temperature. Plants were then held in light at 22 to 24°C at 16 h: 8 h, light: dark cycle, and further after 3 days the cling film was removed. The VIGS phenotypes were observed for up to 29 days. At 24 dpi (days post infiltration), first-pair leaves of infected seedlings along with the mock seedlings were harvested and stored at −80°C for specific transcript analysis.

## Results

### Identification, phylogenetic study, and expression analysis of *OscWRKY1* from *O. sanctum*

To identify the WRKY TF in *O. sanctum*, we use earlier sequence data resources available in the public domain submitted as sequence read archives (SRAs) (Rastogi et al., 2014). From the transcriptome sequence of *O. sanctum*, transcripts that belong to the WRKY superfamily were identified initially. Almost all identified proteins contained highly conserved WRKYGQK motifs in their protein sequences (Fig. S1). After identifying all the *WRKY* transcripts in *O. sanctum*, we BLAST the annotated sequences to identify the full-length transcripts from the transcriptome data. From the total annotated transcripts, the full-length sequence of *OscWRKY1* was amplified using gene-specific primers. It was cloned in the PTZ57R/T vector and further confirmed by sequencing. The size of *OscWRKY1* transcript as per the annotation made in earlier data is 3349 bp, which includes a coding sequence (CDS) of 1023 bp having 340 amino acids. The full-length CDS was aligned with its amino acid encoded sequence as shown in Fig. S2. It was found that the DNA binding domain consists of 58 amino acids. Protein sequences of OscWRKY1 homologs from different plant species were retrieved from the NCBI database for sequence alignment to determine its evolutionary similarity and diversity using Clustal Omega. The phylogenetic homology of OscWRKY1 was determined using MEGAX. OscWRKY1 was showing maximum homology (75%) to the probable WRKY protein of *Salvia splendens* and to the hypothetical protein of *Perilla frutescens* (74%) (Fig. 1a). Based on the BLASTp analysis of WRKY proteins from different species, we found most of them were either putative or uncharacterized (data not shown). From the transcriptome differential gene expression data, we study the expression of unique full-length transcripts present in *O. sanctum* and found that *OscWRKY1, OscWRKY4, OscWRKY48, OscWRKY2,* and *OscWRKY15* were highly expressed (Fig. S3). To further validate the tissue-specific expression patterns of specific full-length *OscWRKY1* genes in leaf and root tissues, we performed real-time PCR. The leaf and root tissue show differential expression of *WRKY* genes; among them, *OscWRKY1’s* expression is higher in leaf tissue than in root tissue (Fig. 1b).

**Figure 1.**
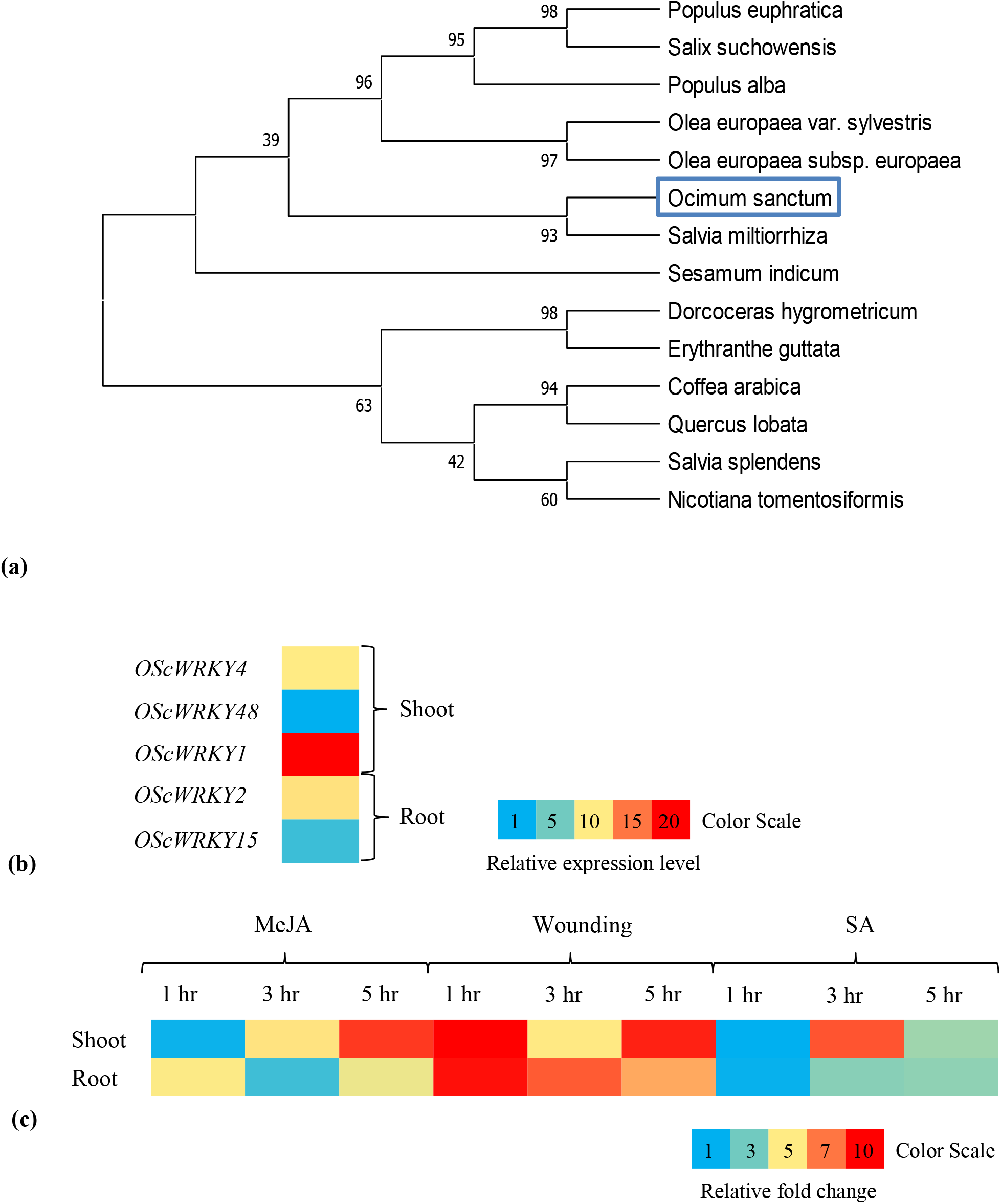
Phylogenetic analysis and tissue specific expression of OscWRKY TF. **(a)** A maximum likelihood phylogenetic tree was constructed using MEGA X software based on the conserved WRKY domains to examine the phylogenetic relationships among different OscWRKYs and it showed the highest homology with *Salvia miltiorrhiza*. The evolutionary history was inferred by using the Maximum Likelihood method and Dayhoff w/freq. model. The bootstrap consensus tree inferred from 1000 replicates is taken to represent the evolutionary history of the taxa analyzed. **(b) Heat map showing tissue-specific expression of *OscWRKY* transcripts.** Some of the selected full-length *OscWRKY* transcripts from the transcriptome data were validated using the qRT-PCR analysis to study the differential expression in the shoot and root tissues. **(c) Tissue-specific expression of *OscWRKY1* after MeJA, SA treatment and wounding.** The relative expression of *OscWRKY1* transcript in the shoot and root tissue was studied after 1, 3, and 5 hours of MeJA, SA treatment and wounding. Relative expression of transcripts was calculated taking untreated plant samples as a control. Actin and ubiquitin were used as an endogenous control for gene normalization. The color scale represents the relative fold change values.

MeJA and SA are known to be important elicitors in diverse signaling cascades and are also involved in the regulation of various enzymes related to plant secondary metabolic pathways. To know the role of identified WRKYs as a signaling molecule in response to MeJA, SA, and wounding, we study the differential expression of *OscWRKY1* after 1, 3, and 5 hours of MeJA, SA treatment, and wounding. We found that *OscWRKY1* is highly induced after 3 and 5 hours of MeJA treatment in the leaf as compared to the root (Fig. 1b). *OscWRKY1* also showed induced expression in response to SA treatment. Wounding in *O. sanctum* also upregulates the early expression of *OscWRKY1* in both leaf and root tissue (Fig. 1c). The significant upregulation of *OscWRKY1* after MeJA, SA, and wound treatment indicates that it might have a role in the regulation of pathogen response or secondary metabolism.

### *OscWRKY1* binds with the promoter containing W-box *cis*-elements and transactivates the reporter gene in yeast

The promoter region contains regulatory components that are required for gene expression to be regulated on a temporal, geographic, and cell type-specific basis (Lescot et al., 2002). For bacterial protein purification and studying the DNA binding affinity, we cloned OscWRKY1 in the pGEX-4T2 expression vector. Recombinant glutathione S-transferase (GST)-OscWRKY1 fusion protein was induced and purified from *E. coli* BL21 (DE3). The size of (GST)-OscWRKY1 is 62.4 KDa. SDS-PAGE was used to check the purified fraction of recombinant OscWRKY1, which was then subjected to an EMSA. The EMSA results indicated that only when OscWRKY1 binds to the W-box *cis*-elements TTGACC (W-box 1) and TTGACT (W-box 5) (Fig. 2a) could DNA-protein complex signals be produced. We designed specific probes for TTGACC and TTGACT *cis*-elements as well as for its mutated counterparts TTGACG and TTCACT with a single nucleotide change. We observed that OscWRKY1 interacts with W-box 1 and W-box 5, while does not able to bind with the mutated probes (Fig. 2b, Fig. S4). These results suggest that W-box 1 and W-box 5 are the novel *cis*-elements bound by OscWRKY1. OscWRKY1 interacts with W-box *cis*-element present within the promoter of target genes with the help of their DNA binding domain (DBD) and thus activates transcription with the help of their transactivation domain. To investigate the transactivation ability, we cloned *OscWRKY1* in pGBKT7 in fusion with modified GAL4-DBD and then transformed into the yeast Y187. OscWRKY1 demonstrated positive transactivation by activating the reporter gene *LacZ* in yeast and producing blue color when X-gal was applied. In the absence of Trp and Ura, transformed yeast colonies were developed in the synthetic dextrose media (Fig. 2c).

**Figure 2.**
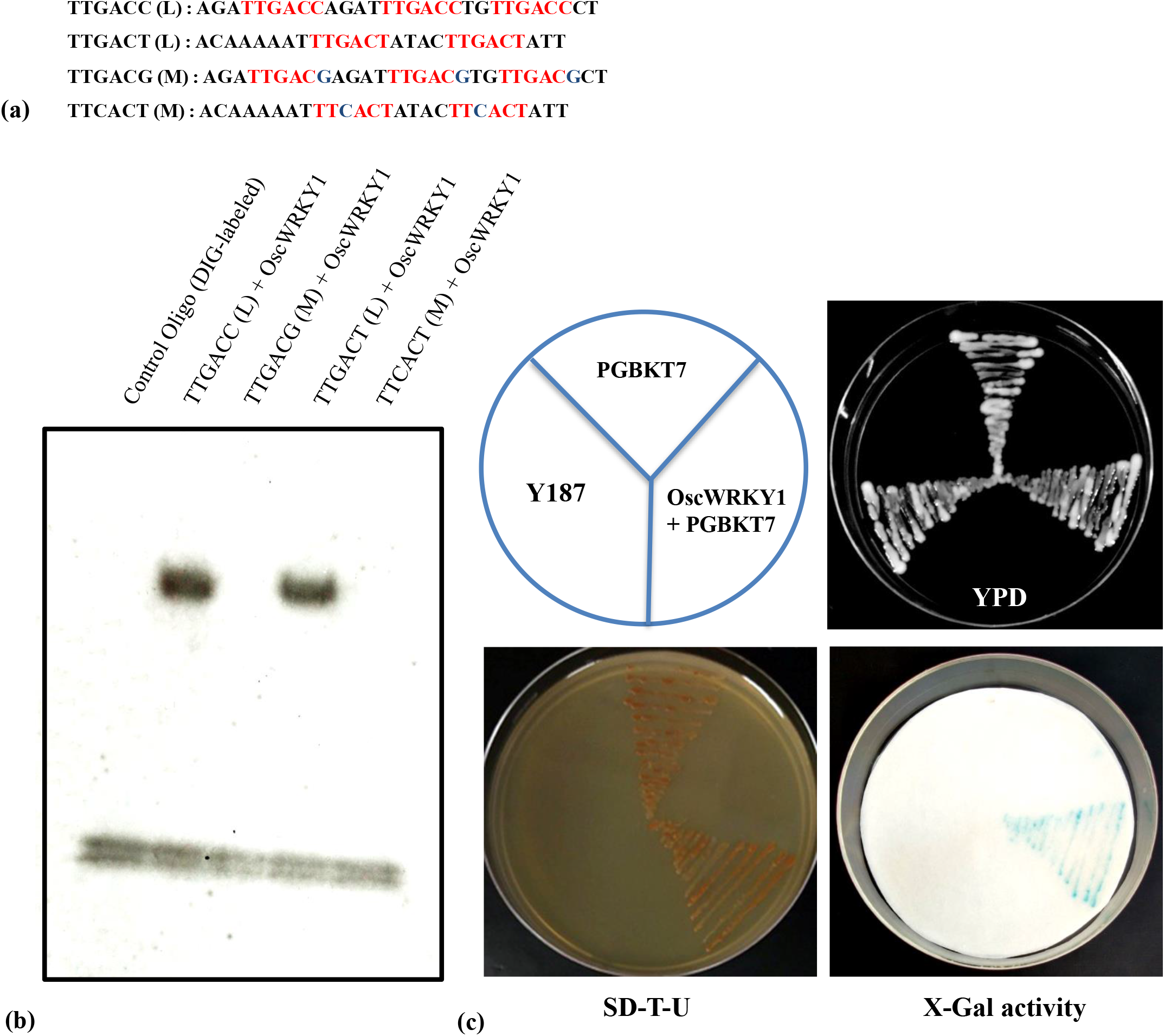
Electrophoretic mobility shift and Transactivation assay of OscWRKY1 protein. **(a)** Probes containing two different W-box *cis*-elements (TTGACC and TTGACT) were designed to study the DNA-protein interaction. The desired *cis*-elements are marked with red color while the binding site carrying mutations are highlighted with blue color. **(b)** EMSA of OscWRKY1 showed that it specifically interacts with both TTGACC and TTGACT cis-elements (L-DIG-labelled and M-mutated probe) M: TTGACG, TTCACT. **(c)** To study the transactivation property of OscWRKY1, it was cloned in yeast expression vector pGBKT-7 in fusion with GAL4-DBD and then transformed in yeast *S. cerevisiae* (Y187 strain). The positive colonies were selected and streaked on dropout SD-U-T media. Transactivation assay was performed using X-gal as a substrate and the resulting blue color showed the positive transactivation property of OscWRKY1 as it could activate the reporter gene *LacZ* in yeast.

### Overexpression of *OscWRKY1* in Arabidopsis confers enhanced resistance to *P. syringae* pv. *tomato* DC3000

To assess the functional role of *OscWRKY1* in plants, transgenic Arabidopsis lines were generated using the constitutive promoter CaMV35S. We selected two independent homozygous lines *(OscWRKY1-L1* and *OscWRKY1-L2)* containing the *35S:OscWRKY1* construct. To validate transgenic lines of Arabidopsis, PCR amplification was performed using genomic DNA and pBI-121 nptII (KanR)/gene-specific primer set (Fig. S5). The predicted amplification product was obtained from transgenic lines but not from WT Col-0 plants, demonstrating that the transgene was successfully incorporated into the Arabidopsis genome. The overexpression of *OscWRKY1* in transgenic plants was confirmed by semi-quantitative PCR using gene-specific primers. We observed that both the homozygous lines *(OscWRKY1-L1* and *OscWRKY1-L2)* have enhanced expression of *OscWRKY1,* whereas no expression was detected in WT Col-0 plants (Fig. S6). To study the role of OscWRKY1 in plant defense, OscWRKY1 overexpressing transgenic plants *(OscWRKY1-L1* and *OscWRKY1-L2)* were infected with bacterial pathogen *Pst* DC3000 and compared with WT Col-0 Arabidopsis plant. Diseased WT Col-0 Arabidopsis leaves displayed severe chlorosis and necrosis at 4 days post-infection (4 dpi). However, at the same stage, transgenic plants overexpressing *OscWRKY1 (OscWRKY1-L1* and *OscWRKY1-L2* lines) exhibited reduced disease symptoms (Fig. 3a). Importantly, estimation of bacterial pathogen *Pst* DC3000 population from the *OscWRKY1-L1* and *OscWRKY1-L2* plants, showed significantly reduced growth of *Pst* DC3000 when compared to WT Col-0 plant (Fig. 3b). At 2 dpi, *Pst* DC3000 growth enhanced by 1.48 [Log (CFU/cm^2^)] unit (~ 30 fold) in WT Col-0 plants, however, during the same time period, *Pst* DC3000 growth enhanced only by 0.58 [Log (CFU/cm^2^)] unit (~ 3.8 fold) and 0.22 [Log (CFU/cm^2^)] unit (~ 1.7 fold), in *OscWRKY1-L1* and *OscWRKY1-L2* plants, respectively (Fig. 3b). Additionally, the Arabidopsis transgenic plants overexpressing *OscWRKY1 (OscWRKY1-L1* and *OscWRKY1-L2)* infiltrated with *Pst* DC3000 showed significantly reduced electrolyte leakage as compared to the WT Col-0 infiltrated with *Pst* DC3000 (Fig. 3c). Furthermore, a GFP-tagged *Pst* DC3000 strain was used to observe the *in planta* colonization of the pathogen in the leaves of WT Col-0, *OscWRKY1-L1* and *OscWRKY1-L2* plants using confocal fluorescence microscopy. The GFP-tagged *Pst* DC3000 bacteria were observed as green fluorescent microcolonies inside the infected leaves. The overexpression transgenic plants *(OscWRKY1-L1* and *OscWRKY1-L2)* displayed reduced colonization bacterial pathogen *Pst* DC3000 as compared to WT Col-0 plants (Fig. 3d). Taken together, these observations suggest that overexpression of *OscWRKY1* enhances the Arabidopsis resistance towards bacterial pathogen, *Pst* DC3000.

**Figure 3.**
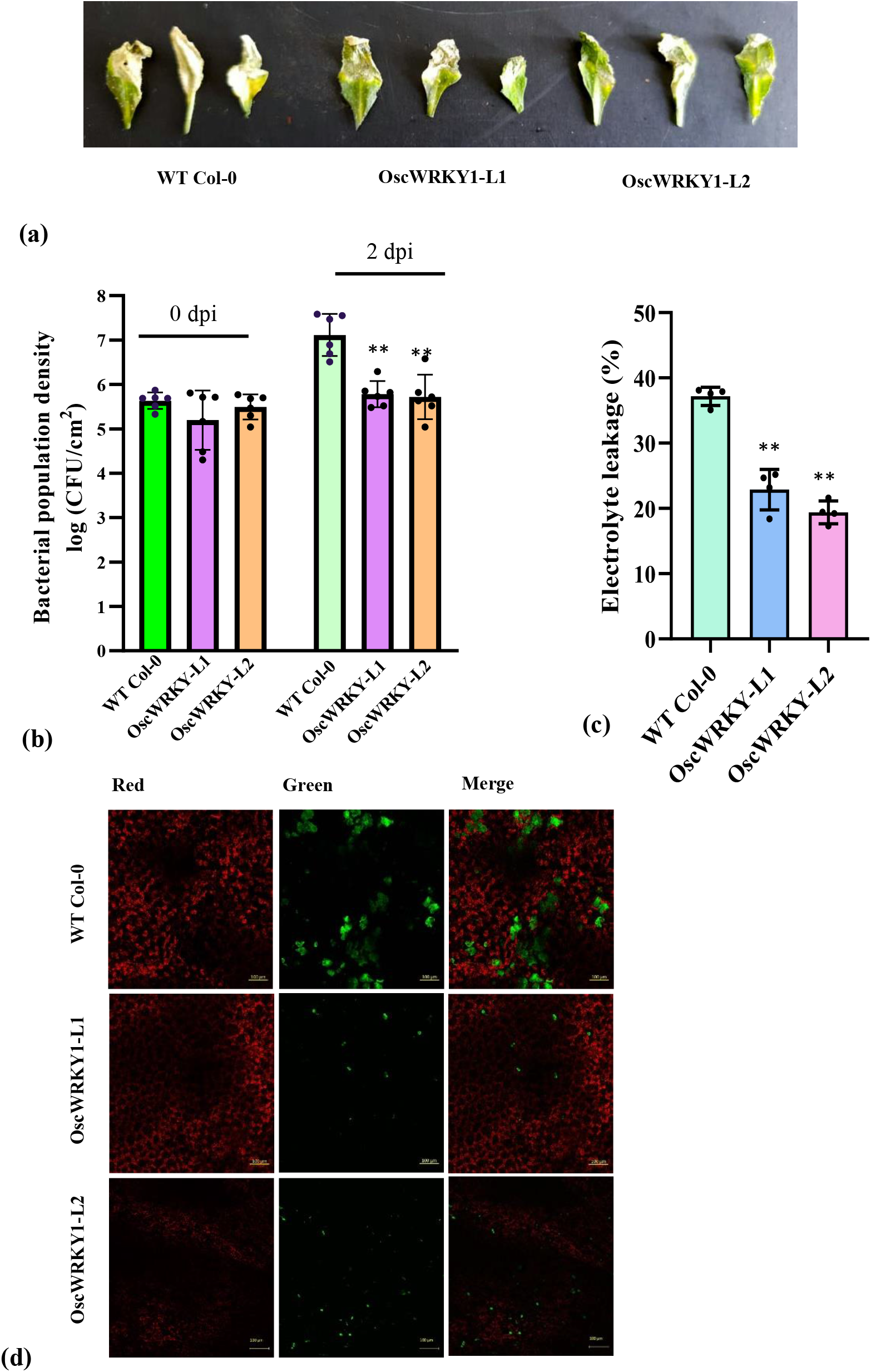
Overexpression of *OscWRKY1* in Arabidopsis promotes resistance against *P. syringae* pv. tomato (*Pst* DC3000). **(a)** Disease phenotype in leaves of WT Col-0, *OscWRKY1-L1* and *OscWRKY1-L2* plants infected with *Pst* DC3000. WT Col-0 and two independent homozygous lines of *Arabidopsis* containing 35S:: *OscWRKY1* construct *(OscWRKY1-L1* and *OscWRKY1-L2)* were grown and leaves of these plants were infiltrated with *Pst* DC3000. Infected leaves were photographed at 4 days post-infection (4 dpi) and are presented in three biological replicates. The overexpression lines of *OscWRKY1* plants display reduced disease symptoms as compared to WT Col-0 plants. **(b)** Estimation of bacterial population in leaves of WT Col-0, *OscWRKY1-L1* and *OscWRKY1-L2* plants infiltrated with *Pst* DC3000 at 2 dpi. The Arabidopsis *OscWRKY1* overexpression lines are more resistant as these plants supported significantly reduced *Pst* DC3000 population as compared to WT Col-0 plant. Data are from two independent experiments and biological triplicates per experiment. Error bars indicate mean ± SD. Student’s t-test: **\***, P < 0.05; **\*\***, P < 0.01. **(c)** Electrolyte leakage was measured in *Arabidopsis* leaves of WT-Col 0, *OscWRKY1-L1* and *OscWRKY1-L2* plants at 2 dpi with *Pst* DC3000. Electrolyte leakage values are presented (%) by calculating the ratio of initial electrical conductivity to the conductivity obtained after autoclaving (100% electrolytes). The leaves of *OscWRKY1* Arabidopsis plants showed lower leakage of electrolytes as compared to wild-type leaves indicating the plant response to the biotic stress. Data are from two independent experiments and biological duplicates per experiment. Error bars indicate mean ± SD. Student’s t-test: **\***, P < 0.05; **\*\***, P < 0.01. **(d)** Visualization of GFP-tagged *Pst* DC3000 in leaves of WT Col-0, *OscWRKY1-L1* and *OscWRKY1-L2* plants using confocal scanning microscopy. The GFP fluorescence was observed with 488 nm excitation and 493-598 nm emission and chlorophyll autofluorescence was observed with 633 nm excitation, and 647-721 emission. Confocal images are shown for GFP fluorescence (green), chlorophyll autofluorescence (red), and merged signals. Scale bars equals 100 μm. Lower colonization of pathogen was detected within the leaves of Arabidopsis overexpression lines infected with GFP-tagged *Pst* DC3000.

### Differential expression study of the phenylpropanoid pathway and PR genes after MeJA treatment and wounding

Increased resistance to plant diseases is generally associated by increased transcript levels of PR genes implicated in the SA defense pathway (Dong et al., 2004; Loake and Grant 2007). Overexpression of *OscWRKY1* increased pathogen resistance in plants; therefore, to further determine whether *OscWRKY1* is involved in controlling the expression of *O. sanctum* phenylpropanoid pathway, we performed qRT-PCR after MeJA and wound treatment in both root and shoot tissue. Transcriptome-wide candidate genes were selected, which are related to the phenylpropanoid pathway and PR genes. Interestingly, the expression of *OscPAL*, *OscC4H,* and *Osc4CL* was induced early after MeJA and wound treatment in leaf tissue (Fig. S7a). PR1 protein II and Thaumatin-like protein were also found to be highly upregulated after 1 hour of MeJA and wound treatment in root tissue (Fig. S7b). These results suggest that *OscWRKY1* elicited by MeJA/wounding might be regulating the expression of phenylpropanoid pathway genes. For further investigation, we analyzed all the promoters of MeJA inducible transcripts in *O. sanctum* and Arabidopsis for the presence of W-box. We found the similar W-box *cis*-elements present in the promoter of Arabidopsis (Fig. S8). *PAL* and *C4H* promoters from both Arabidopsis and *O. sanctum* were amplified from genomic DNA and cloned for further study.

### Interaction study of *OscWRKY1* with the promoter of *AtPAL1* and *AtC4H* from *Arabidopsis thaliana*

In*Arabidopsis thaliana,* overexpression of *OscWRKY1* improved pathogen resistance by interacting with the W-box cis-element present in the promoters of *AtPAL1* and *AtC4H*. The yeast one-hybrid assay is used to confirm the association of *OscWRKY1* with the promoters of *AtPAL1* and *AtC4H*. Reporter constructs for the *AtPAL1* and *AtC4H* promoters were generated by cloning the promoter upstream of the HIS3 reporter gene in the pHIS2.0 vector. To investigate the yeast one-hybrid assay, effector and reporter constructs were co-transformed in *S. cerevisiae* strain Y187. The resulting transformants with *OscWRKY1* and *AtPAL1/C4H* promoters could grow on SD-H-L media. Transformants carrying the *pGAD-OscWRKY1* and *pHIS2.0-AtPAL1/C4H* vectors were unable to grow in the selective media (Fig. 4 a, b). This happens because pGAD-*OscWRKY1* interacts with the W-box cis-element in the promoters of both *AtPAL1* and *AtC4H*, causing transcription of the reporter Histidine (*his*) gene to be activated.

**Figure 4.**
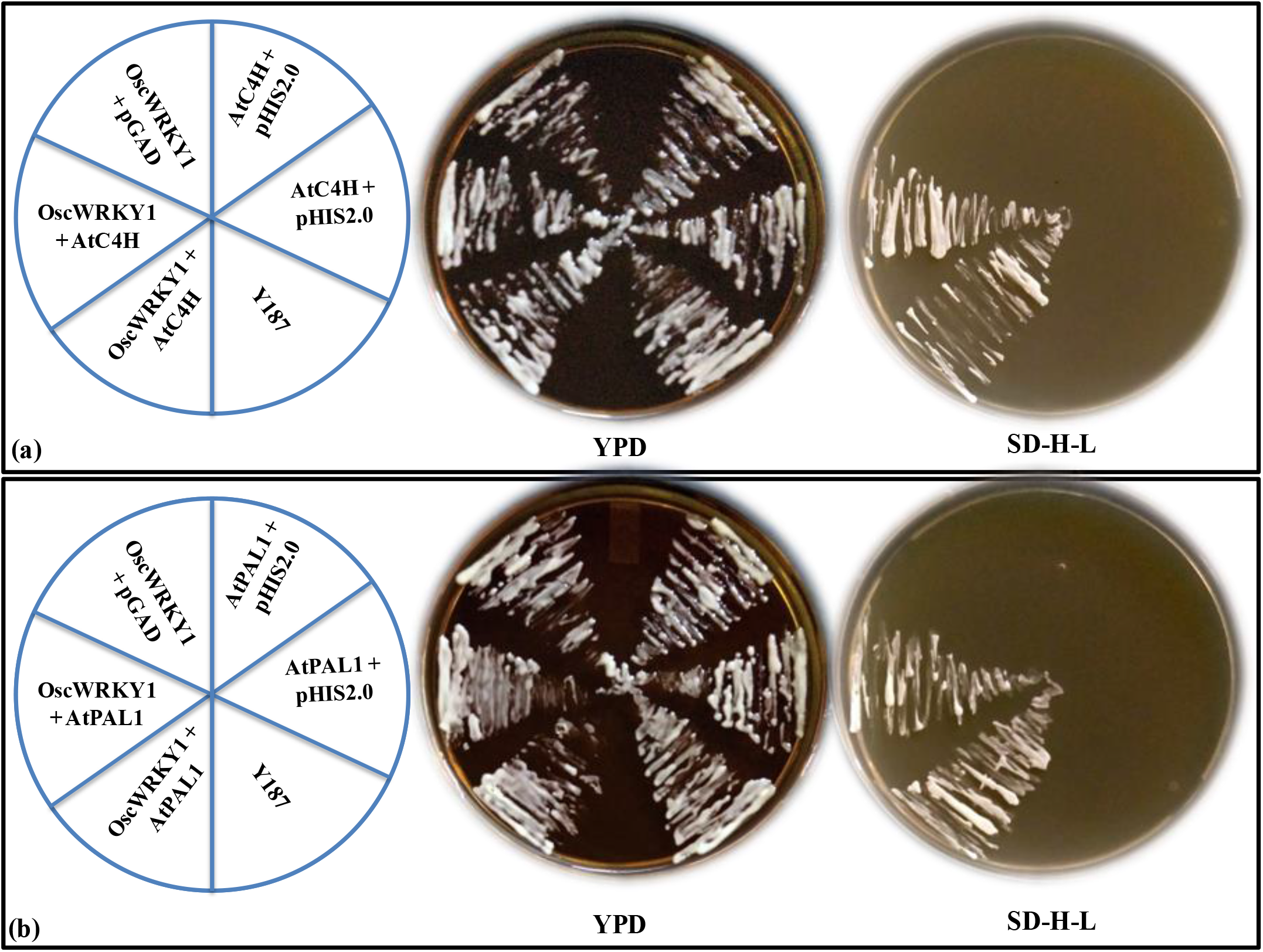
*OscWRKY1* interacts with the promoter of *AtPAL1* and *AtC4H* from *Arabidopsis thaliana.* **(a)** Yeast one-hybrid assay was performed in *S. cerevisiae* (Y187 strain) to study the *in vivo* interaction of *OscWRKY1* with the *AtC4H* promoter. The effector construct was prepared by cloning the *OscWRKY1* transcript in pGAD-T7 vector and the reporter construct was prepared by cloning the desired *AtC4H* promoter containing the W-box cis-elements in pHIS2.0 vector. Both the effector and reporter constructs were then co-transformed in yeast Y187. The positive colonies containing the resulting co-transformants obtained in the selection media were streaked in the YPD and dropout selection media (SD-*his*-*leu*). Positive colonies indicated strong specific interaction of *OscWRKY1* with the *AtC4H* promoter. **(b)** Similarly, the lower panel is the Y1H assay between *OscWRKY1* and *AtPAL1* promoter in yeast Y187. Positive colonies indicated strong specific interactions with the *AtPAL1* promoter. Here the reporter construct was prepared by cloning the desired *AtPAL1* promoter containing the W-box *cis*-elements in pHIS2.0 vector.

### Expression study of *AtPAL1* and *AtC4H* in transgenic *Arabidopsis thaliana*

To further study that *OscWRKY1* regulates the transcript level of *AtPAL1* and *AtC4H* in Arabidopsis, we performed qRT-PCR analysis in control and transgenic lines. We observed an increase in *AtPAL1* expression in both overexpression lines as compared to wild-type Col-0 plants. Additionally, *AtC4H* expression was observed to be elevated in both transgenic lines as compared to WT Col-0 plants. (Fig. 5a). These results suggest the hypothesis that *OscWRKY1* might be regulating *AtPAL1* and *AtC4H* in Arabidopsis by binding with W-box *cis*element within their promoter and involved in increased pathogen resistance in the plant.

**Figure 5.**
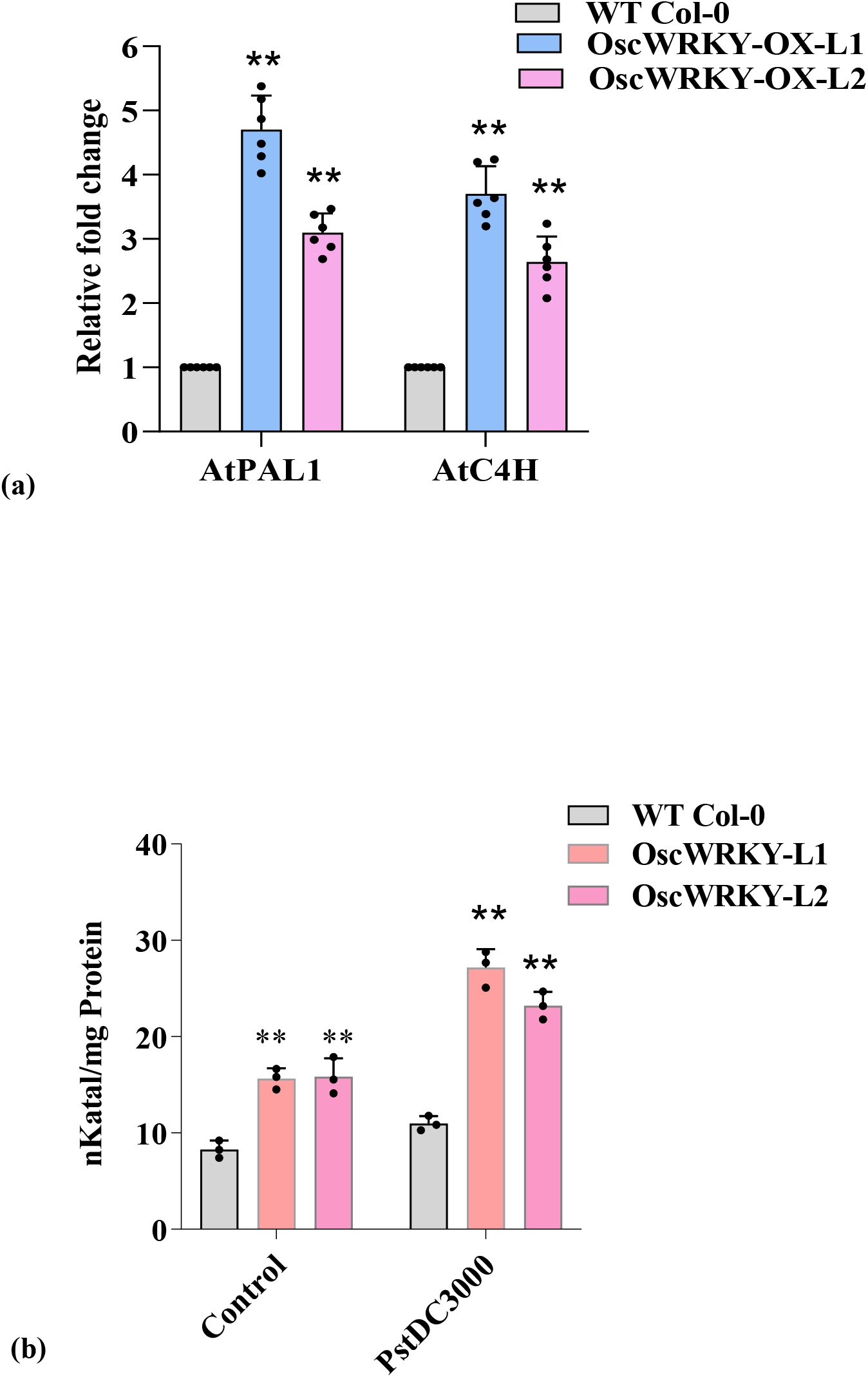
Expression study of PAL, C4H and PAL activity assay in overexpression lines. **(a)** Relative *AtPAL1* and *AtC4H* expression levels were measured through qPCR in the leaves of *OscWRKY1* overexpression lines. The relative expression of *AtPAL1* and *AtC4H* were positively correlated in both *OscWRKY1* overexpression lines. The data represents the run from three independent biological and experimental replicates. Actin, ubiquitin, and eF-1a were used as endogenous control. **(b)** PAL activity is measured in the leaves of *OscWRKY1* overexpression lines and WT Col-0 plants under both control conditions and during infection with *Pst* DC3000. PAL enzyme activity is calculated in nKatal/mg protein. PAL activity is significantly higher in the leaves of *OscWRKY1* transgenic lines. Error bars indicate mean ± SD. Student’s t-test: **\*\***, P < 0.01.

### Enhanced PAL activity in *OscWRKY1* transgenic lines of *Arabidopsis thaliana*

Overexpression study of *OscWRKY1* in Arabidopsis reveals that it might play a crucial role in increased pathogen resistance against *Pst* DC3000. Leaves of WT Col-0 plants showed severe signs of disease symptoms as compared to transgenic *OscWRKY1* plants. Our study shows the binding of *OscWRKY1* with the promoter of *AtPAL1,* which might be involved in regulating PAL content in transgenic plants. We next analyzed PAL activity in both WT Col-0 and *OscWRKY1* overexpression lines under control conditions and during infection with *Pst* DC3000. We observed the level of PAL activity and found that it is significantly higher in the leaves of *OscWRKY1* overexpression lines as compared to WT Col-0 plants under control conditions. As expected, infection with virulent *Pst* DC3000 significantly induces a higher amount of PAL activities in transgenic lines of *OscWRKY1* plants as compared with the WT Col-0 plants (Fig. 5b).

### OscWRKY1 interacts with the promoter of *OscPAL* and *OscC4H* from *O. sanctum*

To validate the DNA binding ability of OscWRKY1 in yeast, the fragment of interest was fused with the GAL4 activation domain present in the pGADT7 vector encoding the leu reporter gene. Reporter constructs for the *OscPAL* and *OscC4H* promoters were constructed by cloning the promoter upstream of the HIS3 reporter gene into the pHIS2.0 vector. Effector and reporter constructs were co-transformed in *S. cerevisiae* strain Y187 to study yeast one-hybrid assay. The resulting transformants containing *OscWRKY1* with *OscPAL/C4H* promoters were unable to grow on SD-H-L media. Transformants containing pGAD-*OscWRKY1* and pHIS2.0-*OscPAL/C4H* vector could not grow on the selective media (Fig. 6 a, b). This occurs as a result of the interaction of pGAD-OscWRKY1 with the W-box *cis*-element in the promoters of both *OscPAL* and *OscC4H,* which activates the reporter Histidine (*His*) gene transcription.

**Figure 6.**
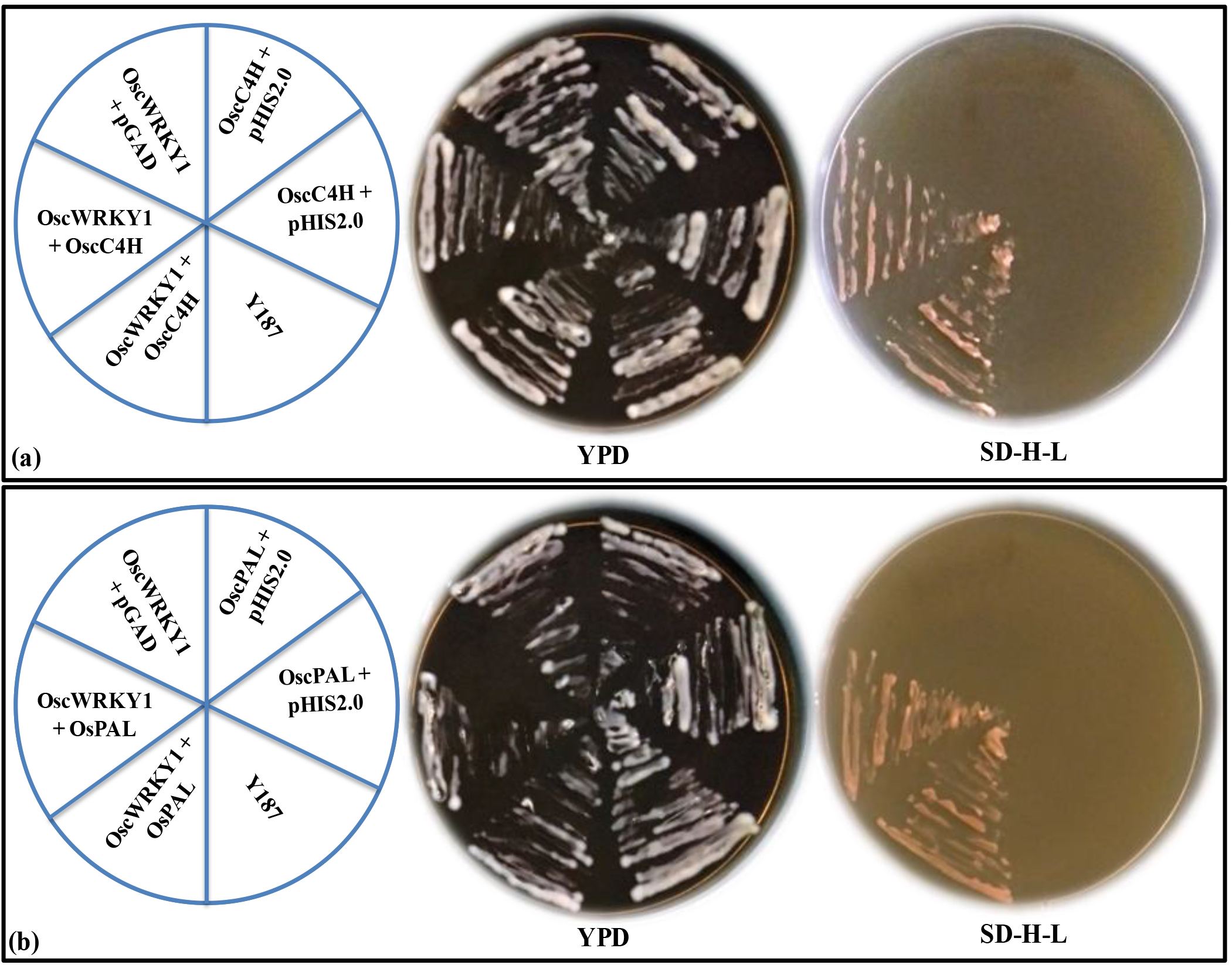
*OscWRKY1* interacts with the promoter of *OscC4H* and *OscPAL.* **(a)** To study the *in vivo* interaction of *OscWRKY1* with the *OscC4H* promoter, Y1H assay was performed in yeast *S. cerevisiae* (Y187 strain). The effector construct was prepared by cloning the *OscWRKY1* transcript in pGAD-T7 vector and the reporter construct was prepared by cloning the *OscC4H* promoter containing the W-box cis-elements in pHIS2.0 vector. The effector and reporter constructs were co-transformed in Y187. The resulting co-transformants obtained in the selection media were streaked in the YPD and dropout selection media (SD-*his*-*leu*). Colonies growing in the selection media showed specific interaction of *OscWRKY1* with the *OscC4H* promoter thus enabling the activation of the reporter gene in yeast. **(b)** Yeast one-hybrid assay between *OscWRKY1* and the *OscPAL* promoter in yeast Y187. Positive colonies indicated strong specific interactions with the *OscPAL* promoter.

### VIGS of *OscWRKY1* confers reduced PAL activity in *O. sanctum*

To assess the efficacy of the silencing vacuum-aided agro-infiltration approach was optimized. *OscPDS* was chosen as a marker gene from *O. sanctum* to ensure the efficiency of the protocol, which leads to the photobleaching of young leaves owing to impaired chlorophyll biosynthesis. The photobleached leaf phenotype as a sign of *OscPDS* silencing started appearing after 13 dpi (days post infiltration) with *Agrobacterium* strains carrying pTRV1 and pTRV2-*OscPDS* vectors (Fig. 7a). However, the symptom gradually became more prominent by 24 dpi. We also study the transcript level of *OscPDS* to understand the effective gene silencing in *O. sanctum* plants. We found that *OscPDS* was downregulated as compared to the vector control (Fig. 7b). To further confirm the silencing of *OscWRKY1,* qRT-PCR was performed in two independent biological replicates.

**Figure 7.**
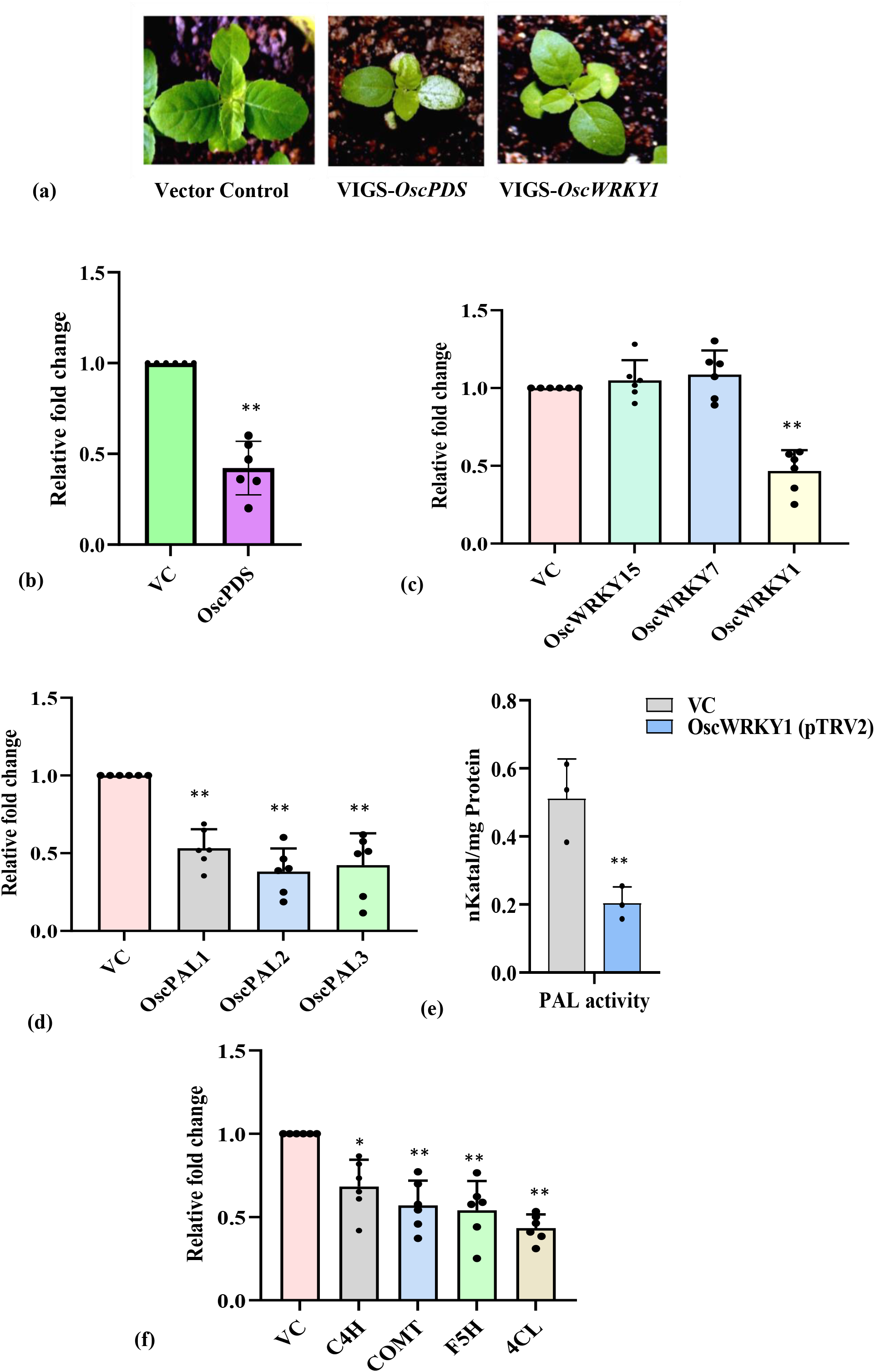
Viral Induced Gene Silencing (VIGS) in *O. sanctum* achieved by vacuum mediated agro-infiltration and expression analysis of phenylpropanoid pathway genes in silenced lines. **(a)** *O. sanctum* plants infected with pTRV1 and pTRV2-*OscPDS*/*OscWRKY1* showing photobleached phenotype 24 days postinfiltration. **(b)** Relative *OscPDS* expression levels in the leaves of empty vector control (pTRV1-pTRV2) and *OscPDS*-pTRV2 infected *O. sanctum* plants based on qRT-PCR analysis. *OscPDS* was used as a marker gene. **(c)** Confirmation of the specificity of silencing by the estimation of the relative transcript level of *OscWRKY1* in vector control and VIGS silenced plants. *VIGS-OscWRKY1* plants showed the downregulation of the *OscWRKY1* gene as compared to the other WRKY genes *(OscWRKY7* and *OscWRKY15)* used as the negative control. **(d)** Relative expression levels of *OscPAL1, OscPAL2* and *OscPAL3* in the leaves of vector control (pTRV1-pTRV2) and *OscWRKY1* -pTRV2 infected *O. sanctum* plants using qRT-PCR analysis. PALs *(OscPAL1, OscPAL2* and *OscPAL3)* were found to be downregulated in silenced lines. The data represent two independent biological and experimental triplicates. Actin, ubiquitin, and eF-1a were used as endogenous control. Error bars indicate mean ± SD. Student’s t-test: **\***, P < 0.05, **\*\***, P < 0.01. **(e)** PAL activity was measured from the leaves of three independent vector control and *OscWRKY1* -pTRV2 plants. Reduced PAL activity was observed in the *OscWRKY1*-pTRV2 plants as compared to the vector control. PAL enzyme activity was calculated in nKatal/mg protein. The data represents error bars with mean ± SD. Student’s t-test: **, P < 0.01. **(f)** Phenylpropanoid pathway gene expression study in vector control and *OscWRKY1* VIGS silenced plants. *VIGS-OscWRKY1* plants showed the downregulation of the *OscC4H, OscCOMT, OscF5H* and *Osc4CL* gene. The data represent two independent biological and experimental triplicates. Actin, ubiquitin, and eF-1a were used as endogenous control. Error bars indicate mean ± SD. Student’s t-test: *, P < 0.05, **, P < 0.01.

We found that *OscWRKY1* was downregulated as compared to the vector control. The expression of some other *WRKY* genes used as a negative control does not show any downregulation in comparison to vector control, which confirms the specificity of VIGS-*OscWRKY1* (Fig. 7c). Additionally, the coat protein (CP) and RNA-dependent RNA polymerase (RdRp) from the pTRV1 and pTRV2 vectors were detected in the agro-infiltrated plants but not in the mock-treated plants. (Fig. S9). This confirms the successful infiltration of pTRV1 and pTRV2 vectors in the seedlings.

Phenylpropanoid biosynthesis is one of the most important processes in secondary metabolism, leading to synthesis of a vast array of natural compounds. *C4H* (Cinnamate-4-hydroxylase), *F5H* (Ferulate-5-hydroxylase), *CCR* (cinnamoyl-CoA reductase), *COMT* (caffeic acid O-methyltransferase) along with the *PAL* (Phenylalanine ammonia lyase) are among the key genes involved in phenylpropanoid biosynthesis. *OscPAL1, OscC4H,* and *Osc4CL* was induced early after MeJA and wound treatment in leaf tissue (Fig. S8a) and *OscWRKY1* was found to interact with the promoters of *PAL* and *C4H* in *Ocimum sanctum* (Fig. 6 a, b). These data imply *OscWRKY1* may regulate phenylpropanoid pathway in *Ocimum sanctum*. To test this theory, we additionally evaluated phenylpropanoid gene expression patterns in *OscWRKY1* VIGS silenced plants*. OscPAL1, OscPAL2* and *OscPAL3* were downregulated in *OscWRKY1* silenced line when compared with the control (Fig. 7d). We next analyzed the PAL activity in both vector control and *OscWRKY1* -pTRV2 agro-infiltrated *O. sanctum* plants and found that PAL activity is reduced in *OscWRKY1*-pTRV2 as compared to the vector control (Fig. 7e). These results further indicate the role of *OscWRKY1* in monitoring the level of PAL activity in the plant by modulating the activity of *OscPAL.* The relative transcript level of *F5H, CCR, COMT, 4CL* and *C4H* were also checked and were found to be downregulated in silenced lines (Fig. 7f).

## Discussion

*O. sanctum* is an important medicinal plant well known for its economically important aromatic oils. It contains aromatic compounds like limonene, eugenol, camphene, α-Pinene, and camphor in their essential oil. *O. sanctum* is rich in phytochemicals, which include mainly terpenoids and phenylpropanoids. Plant WRKY transcription factors are of particular importance as they participate in a variety of biotic and abiotic stress responses and biological processes (Jiang et al., 2015). In this study, we elucidate the functional role of *OscWRKY1* from *O. sanctum* in regulating phenylpropanoid pathway genes. From the earlier published transcriptome data of *O. sanctum*, we have identified the *OscWRKY1* transcripts expressing in the leaf tissue (Rastogi et al., 2014). The phylogenetic study revealed that OscWRKY1 belongs to subgroup III. MeJA and wounding function as elicitors playing a key role in varied plant metabolic processes related to secondary metabolism and defense response (Cheong et al., 2002). The expression of *OscWRKY1* after MeJA, SA, and wounding was found to be higher in comparison with other WRKY transcription factors from *O. sanctum*. We started characterizing *OscWRKY1* due to its higher comparative expression and having a blastX homology with probable WRKY and hypothetical protein from *Salvia splendens* and *Perilla frutescens.*

WRKY genes are known to play a central role in SA- and JA-dependent processes (Thaler et al., 2012). WRKY TF controls gene expression by binding to the W-box *cis*-element, and hence plays an important role in many biological processes. Previous studies revealed the ability of different WRKY TFs to bind with W-box *cis*-elements and their involvement in both positive and negative gene regulation (Wang et al., 2009; Peng et al., 2016; Hu et al., 2018). For example, *WRKY12* negatively regulates cadmium tolerance in *Arabidopsis* by interacting with the promoter of glutathione GSH-1 and represses the expression of phytochelatin biosynthesis genes (Han et al., 2019). *PsWRKY (Papaver somniferum),* and *WsWRKY1 (Withania somnifera)* positively regulates the secondary metabolic pathway genes by binding with the consensus W-box *cis*-element (Mishra et al., 2013; Singh et al., 2017). The present work showed that recombinant OscWRKY1 protein binds with the W-box *cis*-elements (TTGAC[C/T]) and activates the reporter gene in yeast. The yeast one-hybrid experiments further proved that OscWRKY1 binds with the conserved W-box *cis*-elements present in the promoter of *PAL* and *C4H*, which suggests that they might be a direct target of the OscWRKY1 TF.

Plant WRKY TFs acts as both positive and negative regulators, thus exhibiting a crucial role in plant defense response (Pandey and Somssich 2009). In this work, we have studied the biological role of *OscWRKY1* in regulating the phenylpropanoid pathway and plant defense response. The *OscWRKY1* overexpression results in reduced bacterial population density and electrolyte leakage as compared to Col-0 WT Arabidopsis infected with the phytopathogenic bacteria, *Pst* DC3000. Microscopic visualization of Arabidopsis leaves infected with GF-tagged *Pst* DC3000 showed decreased colonization in *OscWRKY1* overexpressing lines as compared to the Col-0 WT. Taken together, these results indicate that the *OscWRKY1* overexpression lines are more resistant towards *Pst* DC3000. In a study, overexpression of jasmonic acid-inducible *WRKY40* from chickpea triggers a defense response against *Pst* in Arabidopsis (Chakraborty et al., 2018). *CaWRKY41,* from *Capsicum annuum* coordinates the responses against pathogenic bacterium *Ralstonia solanacearum* and enhances plant immunity (Dang et al., 2019). Regulation of benzylisoquinoline alkaloids biosynthesis by *StWRKY8* conferred enhanced resistance to late blight disease in potatoes (Yogendra et al., 2017). In a recent study, it has been revealed that differentially regulated *WRKY* genes from roses were specific to biotrophic and a hemibiotrophic leaf pathogen involved in the activation of phenylpropanoid biosynthesis and other factors of the SA signaling pathway (Neu et al., 2019). Phenylpropanoid derivatives can protect plants from biotic infections caused by viruses, bacteria, or fungi. Phenylalanine ammonia lyase knockout mutants in *Arabidopsis* increased susceptibility to a virulent strain of *Pseudomonas syringae* (Huang et al., 2010). Caffeate O-methyltransferase mutants demonstrated reduced resistance to bacterial and fungal infections (Quentin et al., 2009). In certain circumstances, disrupting phenylpropanoid metabolism improved pathogen resistance (Gallego-Giraldo., 2011). External application of a combination of three phenylpropanoids protected plants from *P. humuli* infection (Feiner et al., 2021). *OsWRKY67* from rice directly binds with the promoter of pathogen-related genes *PR1a* and *PR10* and provides bacteria blight resistance (Liu et al., 2018). In cotton, wound responsive *GhWRKY40* plays an important role in enhancing susceptibility to bacterial infection *Ralstonia solanacearum* in transgenic tobacco (Wang et al., 2014).

Due to unavailability of functional mutant in *O. sanctum*, we had utilized VIGS to deliver the more efficient TRV-based VIGS for *O. sanctum.* TRV-based VIGS and its applications have been studied in some medicinal plants, but not in *O. sanctum* (Singh et al., 2015; Liscombe and O’Connor, 2011; Hileman et al., 2005). This is the first report of the VIGS-based gene silencing in *O. sanctum* utilizing *PDS (Phytoene desaturase)* as the marker gene and which further confers the role of OscWRKY1 in regulating the *PAL* gene and thus modulates the PAL activity. Increased PAL activity has been observed during plant-pathogen interactions, however the molecular mechanism of *PAL* activation in response to the pathogen is not known (Li et al., 2015; Abbas et al., 2018; Song et al., 2015; Yu et al., 2016). Overexpression of *CaPAL1* from *Capsicum annum* in Arabidopsis conferred increased pathogen resistance to *Pseudomonas syringae* with increased PAL activity (Kim and Hwang 2014). In rice, *OsMYB30* directly regulates the expression of *OsPALs* in response to Brown plant hopper infestation (He et al., 2020).

We conclude that MeJA/SA and wound responsive TF, *OscWRKY1* binds with the W-box *cis*-elements present within the promoter of phenylpropanoid pathway genes and regulate them in both *O. sanctum* and Arabidopsis. Overexpression of *OscWRKY1* promotes resistance against *Pst* DC3000 in Arabidopsis. Further, *OscWRKY1* silencing using VIGS significantly reduces the five transcripts of phenylpropanoid pathway genes suggesting its major role in regulation of phenylpropanoid pathway.

## Author contributions

AJ has performed the cloning experiments (Arabidopsis promoters, pTRV2, pGADT7 and pHIS2.0), semi-qPCR experiment, Y1H (*O. sanctum* promoter), PAL assay (*O. sanctum* and Arabidopsis), VIGS, *in planta* pathogen colonization experiments, quantification of ion leakage, wrote the manuscript and prepared the figure for the respective part. GSJ has performed the cloning experiments (*Ocimum* promoter, pGEX4T2, pGBKT7, pBI121), qRT-PCR analysis, protein purification, GRA, transactivation assay, Y1H (Arabidopsis promoter), PAL assay (*O. sanctum),* maintained transgenic lines, wrote the manuscript, and prepared the figures. Shikha and Ravi has performed expression analysis of WRKY transcripts from the transcriptome. AP has supervised the pathogenicity assay, bacterial colonization experiments, electrolyte assay and confocal microscopy study. AJ, GJ, AP and RKS have analyzed the result. RKS conceived the theme supervised the work, and edited the manuscript. All authors finally read and approved the manuscript.

## Acknowledgments

The authors acknowledge CSIR-Central Institute of Medicinal and Aromatic Plants (CIMAP) National gene bank for supplying *O. sanctum* seeds. Authors acknowledge Yogesh Kumar and Dr. Feroz Khan for helping in retrieving the genomic sequences of promoters of *O. sanctum*. Authors also acknowledge Payal Srivastava for her help during vacuum infiltration. RKS acknowledges CSIR-CHEMBIO and CSIR-Aroma Mission-I for funding. Gajendra, Ashutosh, and Shikha acknowledge UGC and CSIR for fellowship. Shikha and RKS acknowledge Academy of Scientific and Innovative Research (AcSIR). AP is grateful to DST-INSPIRE Faculty Award (IFA13-LSPA-20).

## Supplementary data

**Fig. S1. Multiple sequence alignment of OscWRKY1.** Almost all identified WRKY proteins from different plant species contained a highly conserved WRKYGQK motif in their protein sequences.

**Fig. S2. Nucleotide and amino acid alignment of *OscWRKY1***. OscWRKY1 contains 1023 bp of ORF, coding 340 amino acids of protein as shown. The amino acid sequence is shown in a single alphabet below each triplet codon. The underlined sequence represents the 58 amino acid long DBD containing the highly conserved WRKYGQK motif shown in the box.

**Fig. S3. Differential gene expression analysis from the transcriptome data of *O. sanctum.*** From the transcriptome differential gene expression data, unique full-length transcripts present in *O. sanctum OscWRKY1, OscWRKY4, OscWRKY2, and OscWRKY15* were found to be highly expressed in leaf and root tissue.

**Fig. S4. EMSA of OscWRKY1.** This figure shows the negative result of OscWRKY1 interaction with the mutated probes.

**Fig. S5. Confirmation of *OscWRKY1* transgenic lines in Arabidopsis.** Genomic DNA PCR of transgenic lines was confirmed by using NptII (KanR) primers, along with an untransformed WT control plant. The expected size of the PCR amplification is 786 bp.

**Fig. S6. Expression analysis of *OscWRKY1* in Arabidopsis transgenic lines.** The overexpression of *OscWRKY1* in transgenic plants was studied by performing semi-quantitative PCR using gene-specific primers and found that both the homozygous lines have high expression of *OscWRKY1,* whereas no expression was detected in wild type Col-0 plants.

**Fig. S7. Differential expression study of the phenylpropanoid pathway and PR genes after MeJA treatment and wounding.** Relative expression of phenylpropanoid pathway genes (a) and PR genes (b) after 1, 3, and 5 hours of MeJA treatment and wounding in both leaf and root tissues was monitored and analyzed. All the samples were carried out in three independent biological (n=3) and experimental replicates. Actin was used as an endogenous control to normalize gene expression. The color scale representing normalized fold induction, as shown in the Figure.

**Fig. S8. Sequence study of PAL and C4H promoters.** PAL and C4H promoter sequences showing the conserved W-box *cis*-element in both *O. sanctum* and *Arabidopsis*. W-box *cis*-elements are highlighted in purple color, and the primer sequences for cloning are highlighted in yellow. The sequence shown with blue color is used for cloning.

**Fig. S9. Determination of efficiency of vacuum mediated agro-infiltration.** Detection of TRV2 coat protein (CP) and TRV1 RNA-dependent RNA polymerase (RdRp) in mock-infiltrated (10 mM MES pH 5.6, 10 mM MgCl_2_, 200 μM acetosyringone), empty vector control (pTRV1-pTRV2) and *pTRV2-OscPDS/OscWRKY1* – infected plants of *O. sanctum* by RT-PCR at 24 days post-infiltration. The presence of CP and RdRP in the leaves of infected plants denotes the successful agro-infiltration of the *Tobacco rattle virus* in *O. sanctum* plants.

**Fig. S10. Melting curve analysis.** After each PCR loop, a melting curve analysis was performed by progressively rising the fluorescence temperature from 60 to 95°C. The qRT-PCR experiments used the 7500 FAST Real-Time PCR system (Applied Biosystems, USA). Actin and ubiquitin were used to normalize gene expression as endogenous controls.

**Table S1.** Primers used in the study.

