## Supplementary figures for "*Ocimum sanctum, OscWRKY1*, regulates phenylpropanoid pathway genes and promotes resistance to pathogen infection in Arabidopsis"

### Figure S1

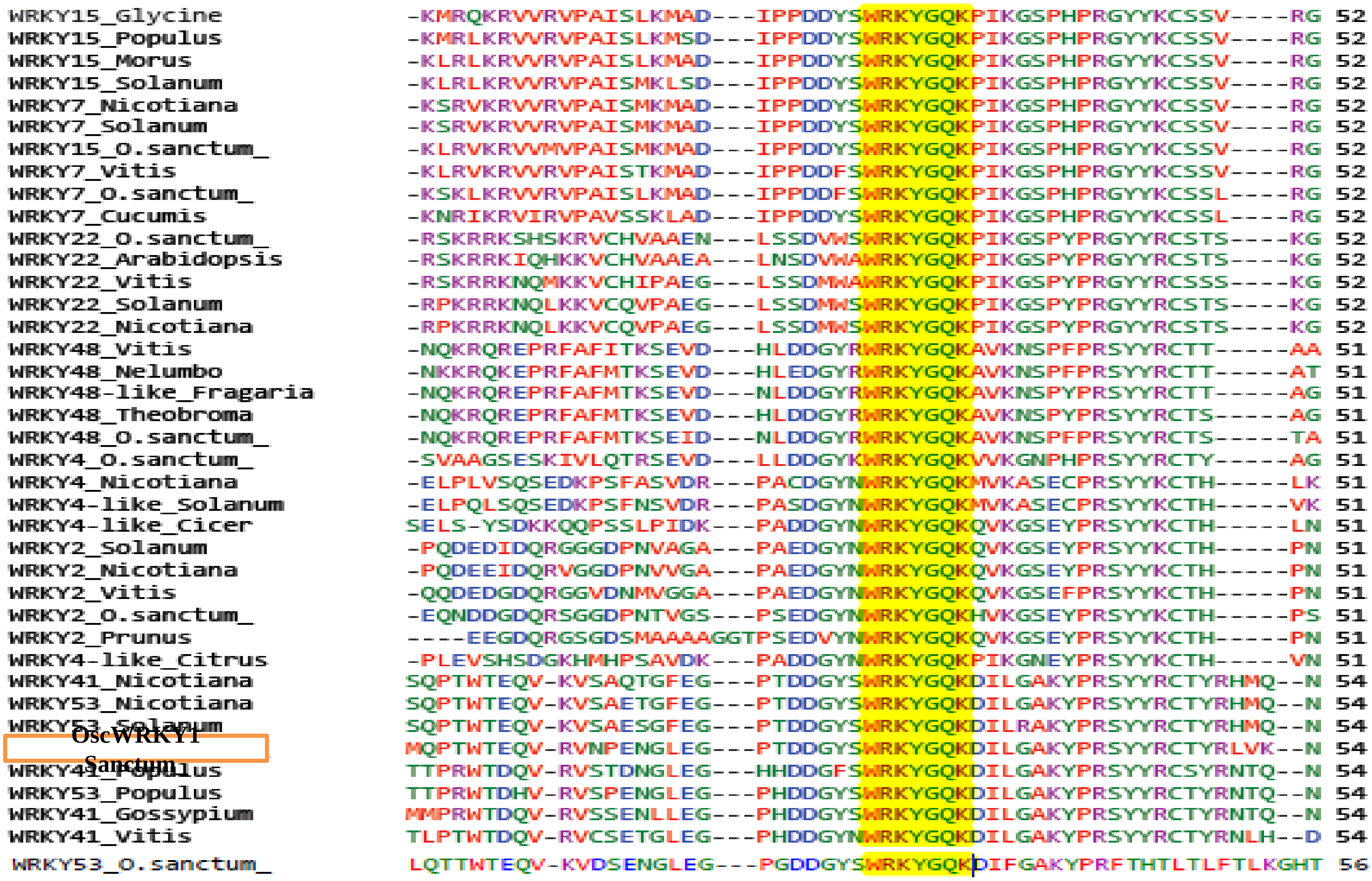


**Fig. S1. Multiple sequence alignment of OscWRKY1.** Almost all identified WRKY proteins from different plant species contained a highly conserved WRKYGQK motif in their protein sequences.

### Figure S2

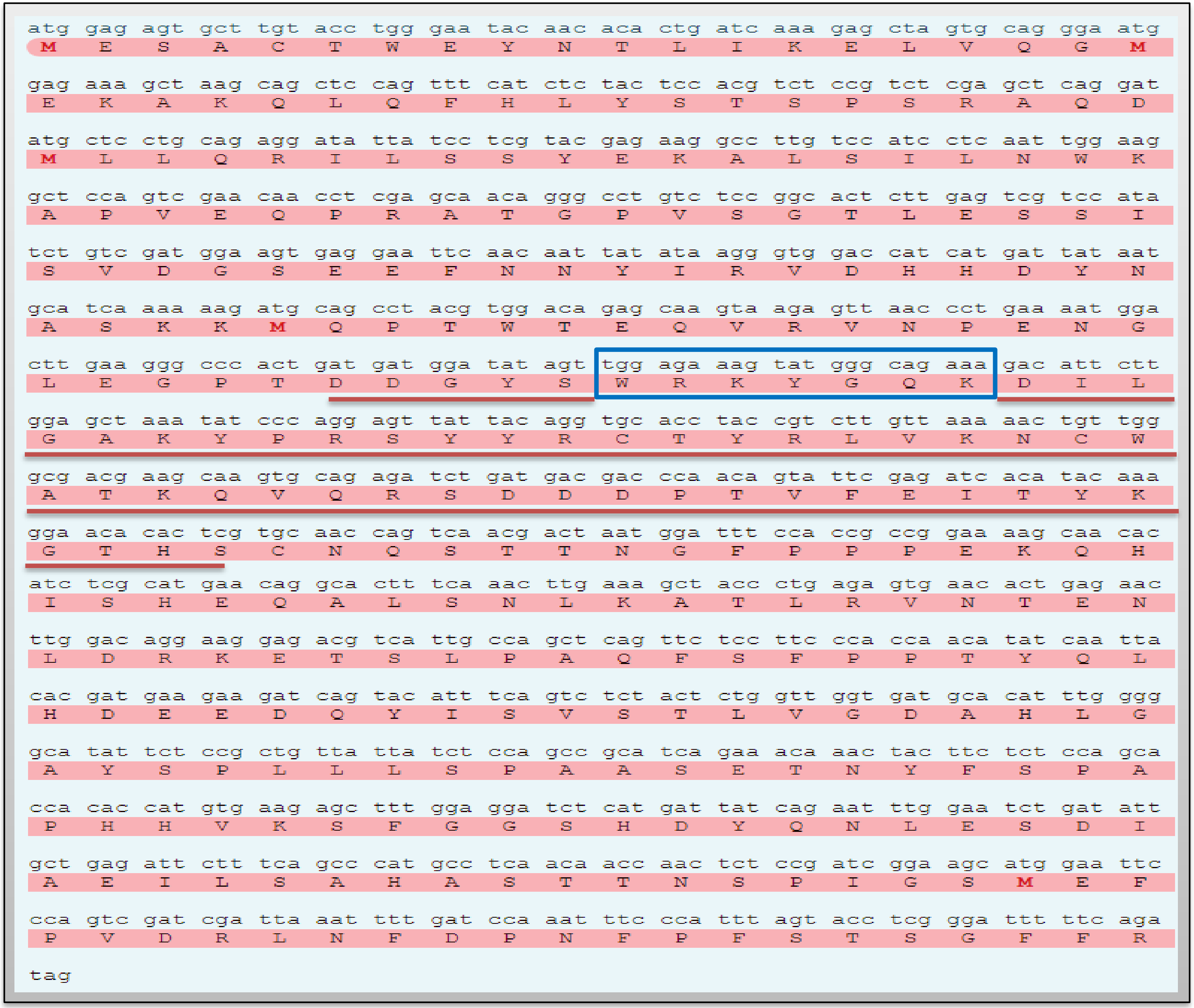


**Fig. S2. Nucleotide and amino acid alignment of *OscWRKY1*.** *OscWRKY1* contains 1023 bp of ORF, coding 340 amino acids of protein as shown. The amino acid sequence is shown in a single alphabet below each triplet codon. The underlined sequence represents the 58 amino acid long DBD containing the highly conserved WRKYGQK motif shown in the box.

**Figure S3**

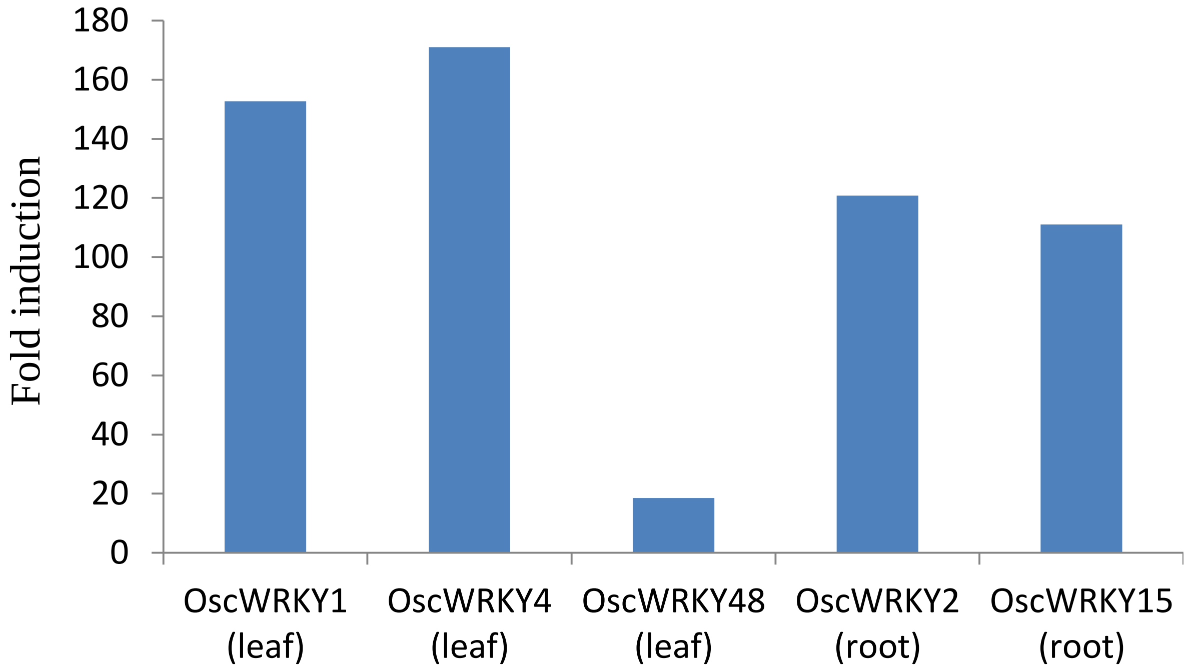


**Fig. S3. Differential gene expression analysis from the transcriptome data of *O. sanctum*.** From the transcriptome differential gene expression data, unique full-length transcripts present in *O. sanctum OscWRKY1, OscWRKY4, OscWRKY2, and OscWRKY15* were found to be highly expressed in leaf and root tissue.

**Figure S4**

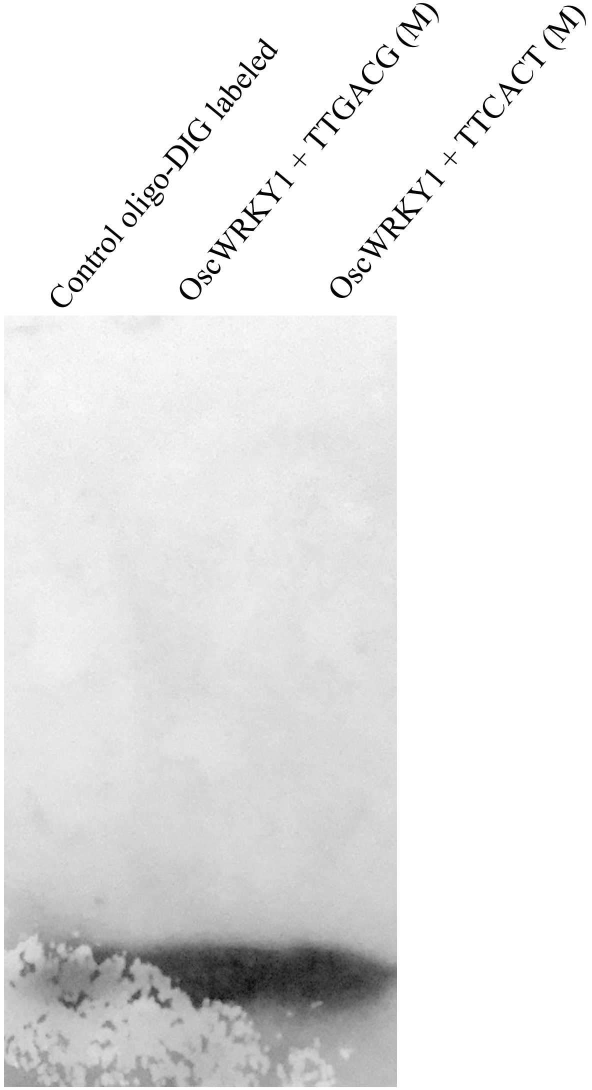


**Fig. S4. EMSA of OscWRKY.** This Figure shows the negative result of OscWRKY1 interaction with the mutated probes.

**Figure S5**

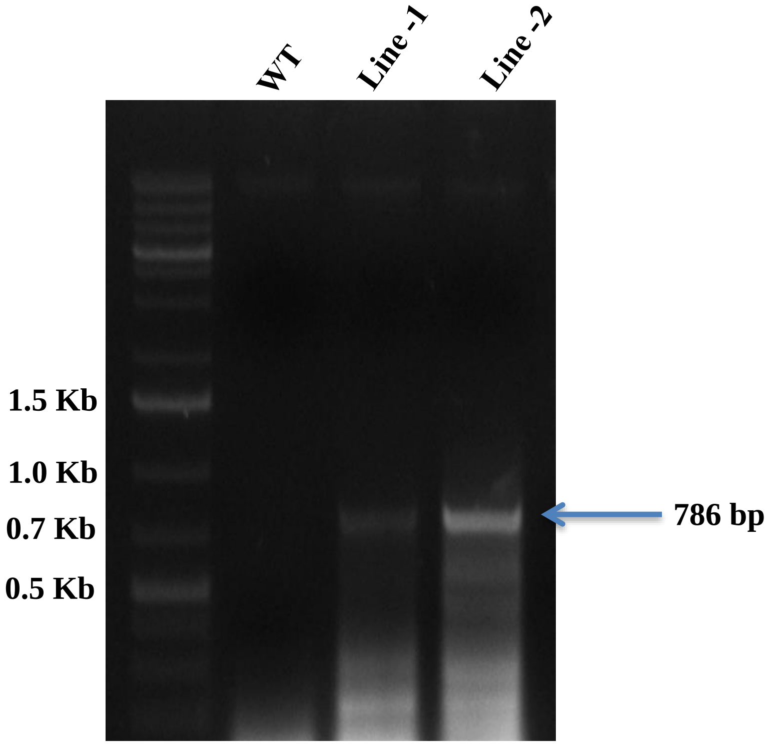


**Fig. S5. Confirmation of *OscWRKY1* transgenic lines in *Arabidopsis.*** Genomic DNA PCR of transgenic lines was confirmed by using NptII (KanR) primers, along with an untransformed WT control plant. The expected size of the PCR amplification is 786 bp.

### Figure S6

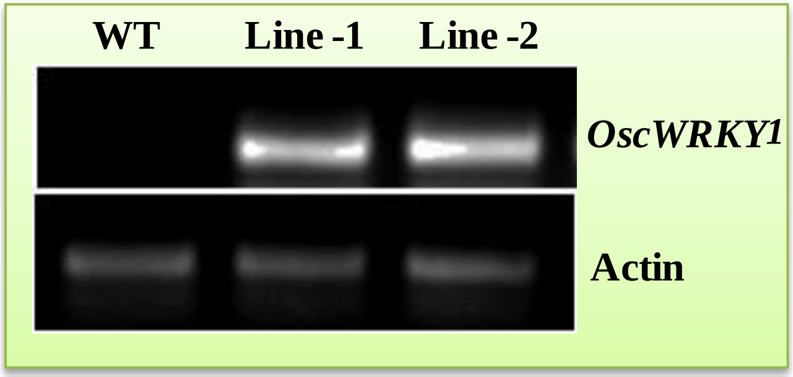


**Fig. S6. Expression analysis of *OscWRKY1* in *Arabidopsis* transgenic lines.** The overexpression of *OscWRKY1* in transgenic plants was studied by performing semi-quantitative PCR using gene-specific primers and found that both the homozygous lines have high expression of *OscWRKY1*, whereas no expression was detected in wild type Col-0 plants.

**Figure S7**

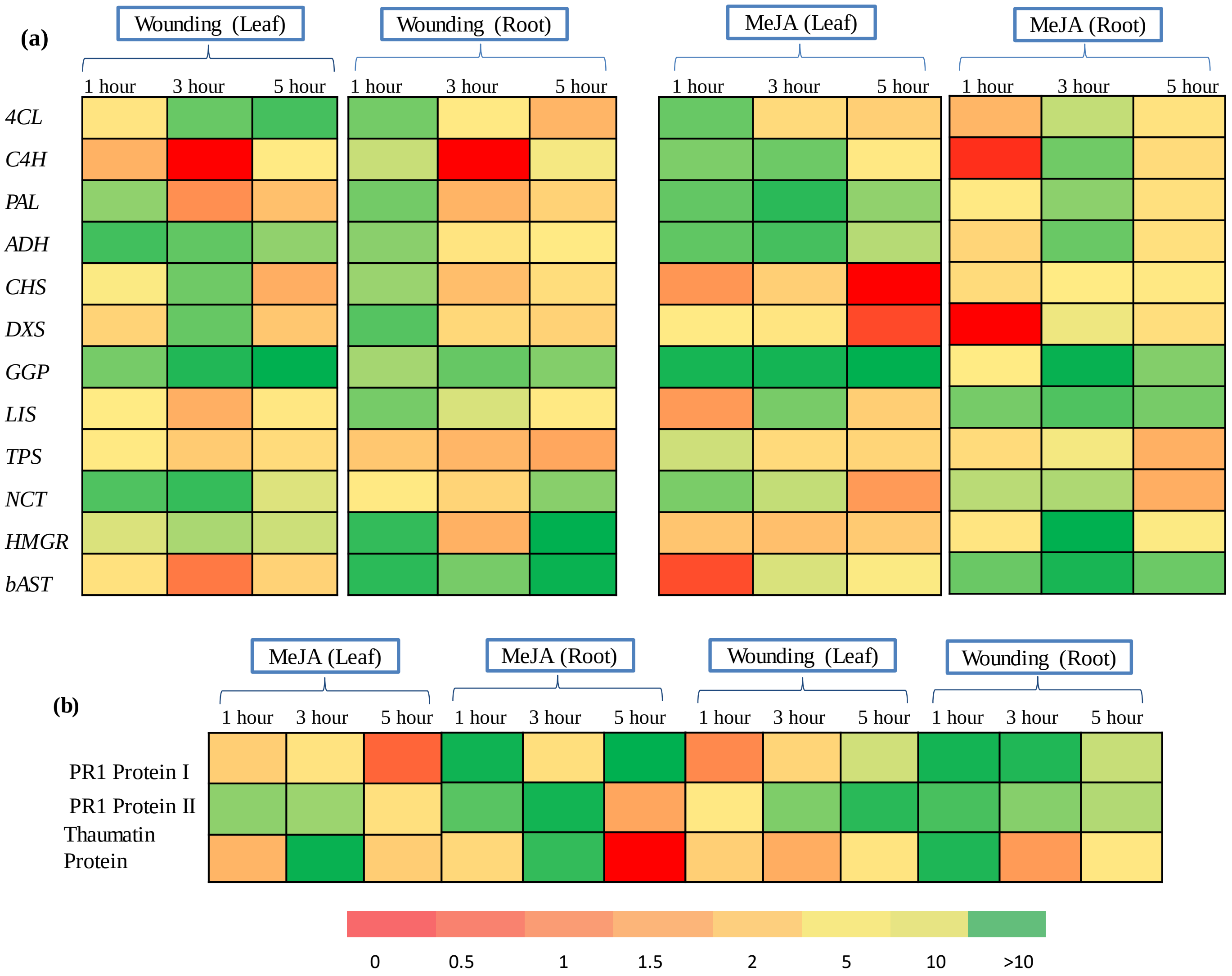


**Fig. S7. Differential expression study of the phenylpropanoid pathway and PR genes after MeJA treatment and wounding.** Relative expression of phenylpropanoid pathway genes (a) and PR genes (b) after 1, 3, and 5 hours of MeJA treatment and wounding in both leaf and root tissues was monitored and analyzed. All the samples were carried out in three independent biological (n=3) and experimental replicates. Actin was used as an endogenous control to normalize gene expression. The color scale representing normalized fold induction, as shown in the Figure.

**Figure S8**

#### PROMOTER SEQUENCE *Osc San*

>Locus_2389_Transcript_4/4_Confidence_0.400_Length_1749 C4H

CCTGTCTAATATTCATCACTACATATTGTACATTCCGTTTGTGTTAATTTCCTACTTTCTCGTATTTGAGCCAGTTCAGTTCAA
CAGTATTTCACCCCGAGTCGCGCTCCGGCGGACCGGTCCCCTTTCGGAGGTCGGGGCGTGACATGGCGATTCAGCTTTCT
ATCTTAATTGTTGACCCACAAACTAACCTAAATCATTTTCAGCTTGCTTTTGTCATCACTCACATGAAATTTTAAGAAACTTT
TCGGAGGATCACTTATCTCAGTTTTACCCTAGACCAAACATGTTCAACTTTGTTATTTTCTTGAAATGAGAATTAAAAAAGG
AATAACTATAAATTATACTCAGTGTTTCTAACATTCCGTAATTTTATCCCAAATTTTGGTTGTCAGGTAAAAATATCTCAAAG
TTCATTTCAACTAGATAAAAATACCCCAAATTGCTTGACGGAGTTATAATGTGCCACGAAGAGACGTTAAATACATGGCGC

>scaffold 945 upstream sequence from -1000 to -1 (Plus/Minus) PAL

CATGGTCCGAATCAGTTACTTAACTATCTATATTTTTAAGTAGATAAAAGGCAAACATTTGCATAACATGGTCTAAAGCATA
CACCTCACCTTTAATTTTTAGGATTTATTTTCAGTTTTTCAACAAAAAAAAAGGGAAATAACACATATTATTAAGCGATATAT
GTATCGTTATAATTGTTATTCTATTGATTAACTTCATCTAATCAACATTACATGACTCAATTTATTTAAATTATGAATGAGGG
TCTAGAATCTTTACTATATTTAATTTGTTCATTCCGTGTATCAATGAATGCAAGTGCAAAAACACGATGGTTTATAATAATA
ATAAAAATGCATAATATTAGATTATTTGGGTGTATATGTGTGGTCAAATAGTATCATAT

#### PROMOTER SEQUENCE *Arabidopsis*

**PAL**

>At2g37040
ATGGTATATATCCTTAATGAGATCCAAGGAAGAGAGAATTAACGTCCCAGACACCTCCAGCACCCACATAATCCATGGACTCTACGTCCACTTT
TAGTCTTTCTATCCATTTTCGTTCGATGGTCGGTTTACTAGTTTTGAAAAAAACCTTTAAAACTTATTTATTGGATAATTATTAAAAACAAAAT
ATAAAATTTATTGGCTCATGATTTTTGCAAAAAAATTCTCTATGTTTTGTTCTCATTTCTCATTTATACGATATTAAAAATTACTTTTCAATAT
GTCGGCCATCATTAAATTGCTTCACTCTAATCCCGTGGCCGTAAACGTTTTTCTTTCAAGCTAGACTCCATAACAACTACGTGCATTTAATTTC
TCAAGTATAAAATTTTCATGTTTTCAAATAGTACGATGTATTTAGTGATTTTATTTATGTACTTTGTTCATTAAATTAGTCATAATTGTTCTGA
TTTTTAGGGGTTTTGATCGAACCCTTAGATCAAAAGTTACCTTAATTGTTTTTTTAGCTAAGTACTTTATTAAAAATTTAATGTTTAGTTCTGA
TTGAGTAGTACTATAAAGGAGACATGTGTCAATCTTGTCAATTGGTTTTGAGTTCAACAATATGCAATATTGCACATGCATTAACGACCAAAAG
AAGATGCAATGCACTTAAATCATTGAAACTGATTTTGTTTTTGTAGTGTATAAAATATCTATTTAATTACCAACGAAAGAAGTGAGCTTTTAAA
AACAAAGAGTCAGAAGATATATATAACTACAAAACCTACAGAAGATAAGCTGGATTTCAAAAGAAGAGAAAGAGTAAACCAATAAATTGACCAA
AGCAAAATCGGATATTTGACATAAGTTTCCATTCACATTGACCCAAATCCACCAGCATTTCAAATAAAGTTACTTAATATAATTTTTGTGTTTA
TAATATATTCCGCCCACTCTTGCCTTCATTTGGACCTTATCCTAAAAGTCAAAACAGGTGAAAAAAATGAGAATACAATTAACACGAAAAATGC
AAAAGACTGTTAAACCGAAATCGAATTCTAGTGTAATCAATCCTTTTCCCAATGATACAACTATAAATCAAAAAGAAAAAATGTACTGATAAAC

**C4H**

>At2g30490
AAAATACGTCACAAATATAATACTAGGCAAATAATTATTTTATTATAAGTCAATAGAGTGGTTGTTGTAAAATTGATTTTTTGATATTGAAAGA
GTTCATGGACGGATGTGTATGCGCCAAATGGTAAGCCCTTGTACTGTGCCGCGCGTATATTTTAACCACCACTAGTTGTTTCTCTTTTTCAAAA
AACACAAAAAAAAAATAATTTGTTTTCTTAACGGCGTCAAATCTGACGGCGTCTCAATACGTTCAATTTTTTTCTTTCTTTCACATGGTTTCTC
ATAGCTTTGCATTGACCATAGGTAAAGGGATAAGGATAATGGTTTTTTCTCTTGTTTGTTTTATCCTTATTATTCAAAAAGGATAAAAAAACAG
TGATATTTAGATTTCTTTGATTAAAAAAGTCATTGAAATTCATATTTGATTTTTTGCTAAATGTCAACACAGAGACACAAACGTAATGCACTGT
CGCCAATATTCATGGATCATGACAATAAATATCACTAGAATAATTAAAAATCAGTAGAATGCAAACAAAGCATTTTCTAAGTAAAACAGTCTTT
TATATTCACGTAATTGGAATTTCCTTTTTTTTTTTTTGTCGTAATTGGAATTTCCTTTATCAAACCCAAAGTCCAAAACAATCGGCAATGTTTT
GCAAAATGTTCAAAACTATTGGCGGGTTGGTCTATCCGAATTGAAGATCTTTTCTCCATATGATAGACCAACGAAATTCGGCATACGTGTTTTT
TTTTTGTTTTGAAAACCCTTTAAACAACCTTAATTCAAAATACTAATGTAACTTTATTGAACGTGCATCTAAAAATTTTGAACTTTGCTTTTG
AGAAATAATCAATGTACCAATAAAGAAGATGTAGTACATACATTATAATTAAATACAAAAAAGGAATCACCATATAGTACATGGTAGACAATGA
AAAACTTTAAAACATATACAATCAATAATACTCTTTGTGCATAACTTTTTTTGTCGTCTCGAGTTTATATTTGAGTACTTATACAAACTATTAG

**Fig. S8. Sequence study of PAL and C4H promoters.** PAL and C4H promoter sequences showing the conserved W-box *cis*-element in both *O. sanctum* and *Arabidopsis*. W-box *cis*-elements are highlighted in purple color, and the primer sequences for cloning are highlighted in yellow. The sequence shown with blue color is used for cloning.

**Figure S9**

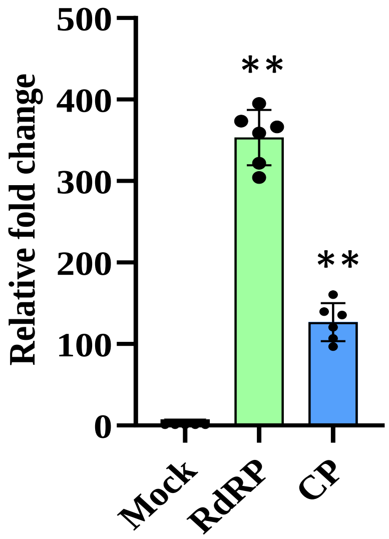


**Fig. S9. Determination of efficiency of vacuum mediated agro-infiltration.** Detection of TRV2 coat protein (CP) and TRV1 RNA-dependent RNA polymerase (RdRp) in mock-infiltrated (10 mM MES pH 5.6, 10 mM $MgCl_2$, 200 µM acetosyringone), empty vector control (pTRV1-pTRV2) and pTRV2-*OscPDS*/*OscWRKY1* -infected plants of *O.sanctum* by RT-PCR at 24 days post-infiltration. The presence of CP and RdRP in the leaves of infected plants denotes the successful agro-infiltration of the *Tobacco rattle virus* in *O. sanctum* plants.

**Figure S10**

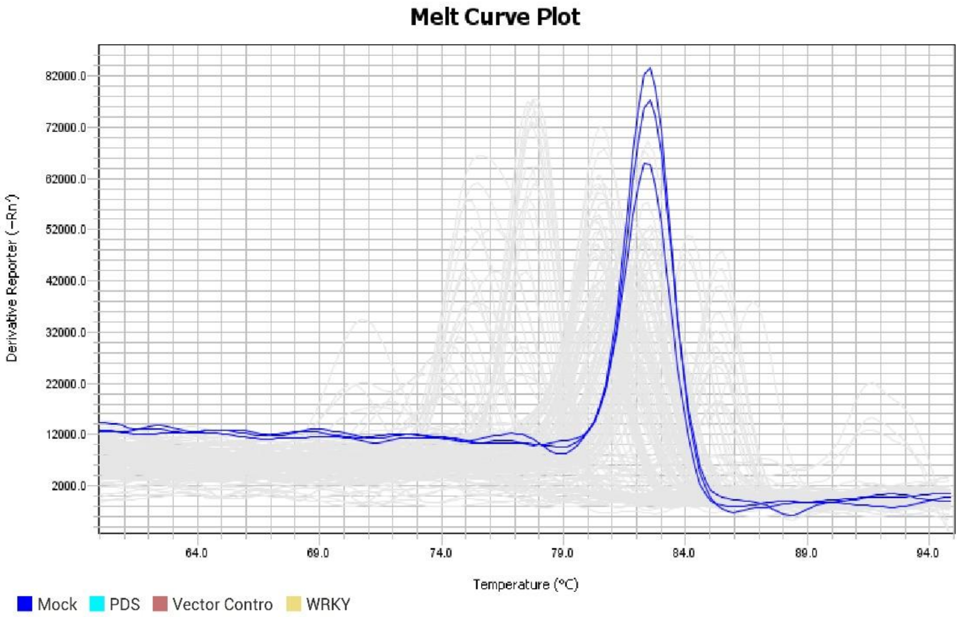


**Fig. S11. Melting curve analysis.** After each PCR loop, a melting curve analysis was performed by progressively rising the fluorescence temperature from 60 to 95°C. The qRT-PCR experiments used the 7500 FAST Real-Time PCR system (Applied Biosystems, USA). Actin and ubiquitin were used to normalize gene expression as endogenous controls.
